# Lung hypoperfusion stimulates liquid absorption in alveoli

**DOI:** 10.64898/2026.06.29.735362

**Authors:** Jimmy Zhang, Deebly Chavez, Sayahi Suthakaran, Chaya Sussman, Stephanie Tang, Sarah K. L. Moore, Clemente J. Britto, Jaymin J. Kathiriya, Hooman D. Poor, Jaime L. Hook

**Affiliations:** Division of Pulmonary, Critical Care and Sleep Medicine, Department of Medicine, Icahn School of Medicine at Mount Sinai, New York, NY, USA; Graduate School of Biomedical Sciences, Icahn School of Medicine at Mount Sinai, New York, NY, USA; Division of Pulmonary, Critical Care, and Sleep Medicine, Department of Medicine, Yale University School of Medicine, New Haven, CT, USA; Department of Stem Cell Biology and Regenerative Medicine, Icahn School of Medicine at Mount Sinai, New York, NY, USA; Mount Sinai Respiratory Institute, Mount Sinai Health System, New York, NY, USA; Department of Microbiology, Icahn School of Medicine at Mount Sinai, New York, NY, USA; Global Health and Emerging Pathogens Institute, Icahn School of Medicine at Mount Sinai, New York, NY, USA

## Abstract

Tissue hypoperfusion is common in clinical settings, but how tissues respond to hypoperfusion on a microphysiological scale is not clear. We used real-time confocal microscopy of live, perfused lungs to gain insights into the effects of hypoperfusion on the microcirculation and microphysiology of lung alveoli, where gas exchange occurs. We focused on effects of hypoperfusion on alveolar liquid secretion, since alveolar liquid secretion is important for alveolar homeostatic functions. Our findings show lung hypoperfusion stimulated a reversal of alveolar liquid transport, from homeostatic liquid secretion to absorption. Specifically, lung perfusion at or near physiological perfusion pressure led to alveolar liquid secretion that depended on the alveolar epithelial cystic fibrosis transmembrane conductance regulator (CFTR), Na^+^-K^+^-Cl^-^ cotransporters, and the Na^+^/K^+^-ATPase. Within minutes of halting lung perfusion or majorly reducing it, alveoli stopped secreting liquid and instead absorbed it via the epithelial Na^+^ channel, CFTR, and K^+^-Cl^-^ cotransporters. We provide evidence that hypoperfusion caused alveolar microvessel lumens to shrink and airspaces to expand, leading to epithelial stretch that stimulated liquid absorption. These findings show lung hypoperfusion initiates mechanical signals that stimulate the alveolar epithelium to absorb liquid, and they may inform the pathogenesis of lung diseases characterized by acute microvascular hypoperfusion.

## INTRODUCTION

Tissue hypoperfusion occurs often in acute medical and surgical conditions that affect the circulatory system. Its clinical impact is substantial, since it associates with organ failure and mortality in conditions as diverse as hemorrhagic shock, transplantation, sepsis, pneumonia, heart failure, and heart surgery (1–6). Tissue responses to acute hypoperfusion are not yet clear, but gaining a better understanding of them might lead to improvements in the clinical care of the critically ill (7). Here, we considered how the lung responds to hypoperfusion. Our rationale for focusing on the lung is that reports suggest the lung is uniquely susceptible to hypoperfusion-induced tissue injury (8), and lung tissue injury has the potential to impact extrapulmonary organs through hypoxemia and other pathways.

The current understanding of lung responses to hypoperfusion focuses largely on lung macrovascular hemodynamics, lung inflammation, and pulmonary edema. Early studies in dogs show blood loss-induced decrease of mean arterial pressure causes a nearly immediate decrease of pulmonary artery pressure (PAP) and increase of pulmonary vascular resistance (9–11), indicating the lung experiences, senses, and rapidly responds to systemic hypotension. Stopping, then restarting lung blood flow reveals a similar increase of pulmonary vascular resistance (12, 13), indicating that lung responses to hypoperfusion survive transient lung ischemia. These macrovascular responses are universally accompanied by lung inflammation (10, 13, 14). However, while stopping lung blood flow for hours before restarting it reliably causes airspace edema formation, systemic hypotension does not (10–17), suggesting that only lung hypoperfusion in its most extreme form – that is, the prolonged absence of blood flow – causes alveolar damage. It remains unclear how alveoli are affected by the less extreme lung hypoperfusion events that are probably more common in clinical settings.

Reports of alveolar responses to lung hypoperfusion are scarce. One line of evidence shows reducing lung perfusion pressure causes alveolar microvascular blood volumes to decrease and lumens to contract (18, 19), indicating that lung hypoperfusion alters alveolar morphology. A second line of evidence shows lung hypoperfusion leads to decrease of microvascular lumen size at flat septa of alveoli but not at alveolar corners (20), indicating that hypoperfusion-induced morphological changes in alveoli are spatially heterogeneous. These data provide important insights into how lung hypoperfusion affects alveolar structure. Yet, an understanding of how lung hypoperfusion affects alveolar function remains elusive. Reports that show the distal lung and cultured alveolar epithelial cells respond to *increases* of hydrostatic pressure with changes of ion and liquid transport (21–23) suggest that *decreases* of hydrostatic pressure in hypoperfused lungs might similarly disrupt alveolar liquid dynamics. Such changes could have implications for lung health, since liquid transport in alveoli maintains the epithelial surface lining layer and contributes to host defense (24).

In this report, we present evidence of how alveolar function changes in response to lung hypoperfusion. Our findings show lung hypoperfusion stimulated a reversal of liquid transport in alveoli of live, intact mouse lungs. Specifically, lung perfusion at or near physiological levels led to alveolar liquid secretion, while halting lung perfusion or majorly reducing it stimulated alveolar liquid absorption. The homeostatic liquid secretion depended on the alveolar epithelial cystic fibrosis transmembrane conductance regulator (CFTR), Na^+^-K^+^-Cl^-^ cotransporters (NKCCs), and the Na^+^/K^+^-ATPase, while the hypoperfusion-induced liquid absorption depended on the epithelial Na^+^ channel (ENaC), CFTR, and K^+^-Cl^-^ cotransporters (KCC) and was accompanied by inhibition of surfactant secretion. We provide evidence that hypoperfusion-induced contraction of alveolar microvascular lumens and expansion of airspaces caused alveolar epithelial stretch that stimulated the switch from liquid secretion to absorption. Together, these data show alveoli dynamically regulate homeostatic functions in response to perfusion-related mechanical cues, such that lung hypoperfusion rapidly stimulates alveoli to absorb liquid and halt surfactant secretion. Loss of homeostatic liquid and surfactant secretion in alveoli of hypoperfused lungs might have a role in the pathogenesis of lung diseases characterized by acute microvascular hypoperfusion.

## RESULTS

### Lung hypoperfusion stimulates a switch from alveolar liquid secretion to absorption

We applied our established methods (25, 26) to excise, inflate, perfuse, and image live mouse lungs by upright confocal microscopy (**Figure 1, A-B**). Contents of the lung perfusate solution are shown in **Supplemental Table 1**. As we have done previously (25, 26), we established lung perfusion using a roller pump that generated liquid flow in a tubing circuit that was connected to the live lungs via two cannulas: one that entered the heart in the right ventricle, crossed the pulmonic valve, and terminated in the pulmonary artery (PA); and one that entered the heart in the left ventricle, crossed the mitral valve, and terminated in the left atrium (LA) (**Figure 1, A-B**). Since mean PA pressure in mice ranges 10-18 mm Hg (27–29) or 13-25 cm H_2_O, we initiated lung perfusion at PAP 15 cm H_2_O as measured by an in-line pressure transducer positioned at the PA cannula and at the same height as the imaged surface of the lung (**Figure 1B** and **Supplemental Table 2**). In separate experiments, we decreased the roller pump speed to achieve PAP 10 cm H_2_O or 5 cm H_2_O. We stopped perfusion in some lungs by halting the roller pump. Left atrial pressure (LAP), which was the pressure in an in-line pressure transducer positioned at the LA cannula and at the same height as the imaged surface of the lung, was 0 cm H_2_O when the roller pump was halted and 1-4 cm H_2_O when the PAP was 5-15 cm H_2_O (**Supplemental Table 2**). LAP decreased concomitant with PAP decreases (**Supplemental Table 2**), consistent with published models of lung hypoperfusion (15, 16). For the remainder of this report, we refer to the experimental conditions as “perfusion at PAP 15 cm H_2_O”, and so on.

**Figure 1.**
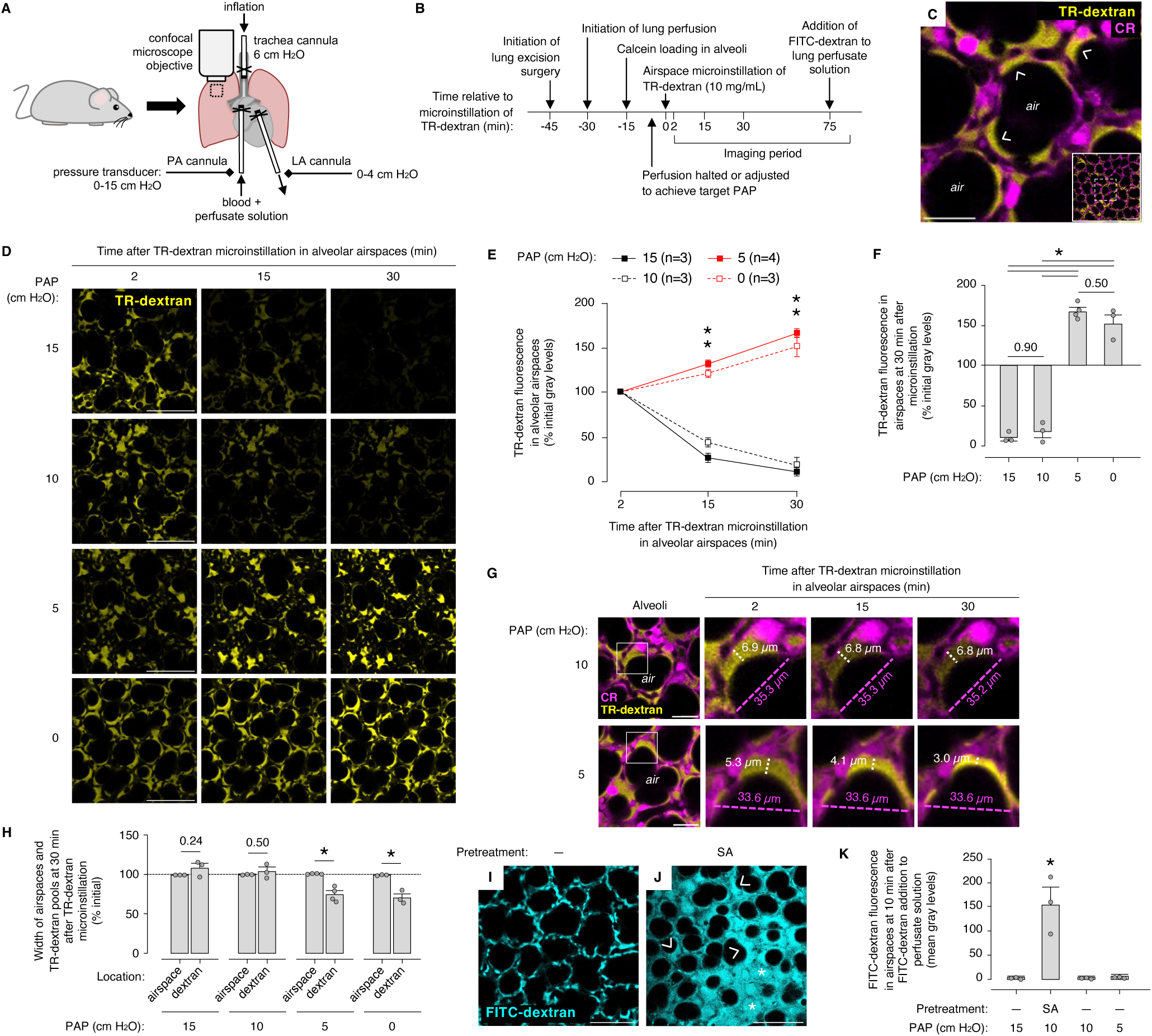
Lung hypoperfusion stimulates a switch from homeostatic AWL secretion to airspace liquid absorption. (**A-K**) Cartoons in <u>A-B</u> show the design of confocal imaging experiments in live, perfused lungs. Determinations of alveolar liquid transport were performed using TRITC-labeled dextran (*TR-dextran*, 70 kDa). See text. *PA,* pulmonary artery; *LA,* left atrium. Images in <u>C</u> show low (*inset*) and high power views of TR-dextran fluorescence (*yellow*) in alveoli. Fluorescence of calcein red-orange (*CR*, *magenta*) delineates alveolar walls; *dashed square* shows the location of the high power view; *arrowheads* show TR-dextran that pooled at alveolar niches. *Air,* example airspace. Images and group data in <u>D-F</u> show change of alveolar TR-dextran fluorescence in lungs perfused at the indicated pulmonary artery pressure (*PAP*). CR fluorescence is not shown. In E and F, n (E) and *circles* (F) each represent mice in which mean fluorescence (E) or mean fluorescence change (F) was quantified in imaging fields of at least 50 alveoli. *Bars* in F show mean ± SEM. \**p* < 0.05 or as indicated by two-tailed *t* test versus PAP 15 cm H_2_O (E) or by ANOVA with post hoc Tukey testing versus *first bar* (F). Images and group data in <u>G-H</u> show change of TR-dextran pool width (*white line and text*) and airspace width (*magenta lines and text*) over time. The *squares* in column 1 in G show the locations of the high power views in columns 2-4. In H, *circles* and *bars* are as in F, but \**p* < 0.05 or as indicated by two-tailed *t* test. Images and group data in <u>I-K</u> show fluorescence of FITC-labeled dextran (20 kDa, 5 mg/mL, *cyan*) added to the lung perfusate solution as indicated in A. Some mice were pretreated with intranasal instillation of *S. aureus* (*SA*) 6 h before live lung imaging, as indicated. *Asterisks* and *arrowheads* point out airspaces that are completely and partially filled with FITC-dextran, respectively. In K, *circles, bars,* and *p* values are as in F. Scale bars: 30 (C,G) and 100 (C *inset* and D,I,J) μm.

We visualized real-time liquid transport in live alveoli of the perfused lungs using an established method in which fluorophore-tagged dextran in alveolar airspaces serves as an indicator of airspace liquid secretion and absorption (26, 30). To apply this method, we first micropunctured single alveoli under bright field microscopy to microinstill alveolar airspaces with the viability dye, calcein red-orange. Calcein fluorescence in the alveolar epithelium delineated alveolar walls and indicated the alveolar epithelium was viable (**Figure 1C**). Airspace fluorescence of microinstilled, fluorophore-conjugated anti-CD11c antibody identified alveolar macrophages (**Supplemental Figure 1, A-B**), affirming that expected non-epithelial cell types were present in alveoli of intact, perfused lungs. Next, we microinstilled alveolar airspaces with a 2-second microinfusion of tetramethylrhodamine isothiocyanate-conjugated dextran in aqueous vehicle solution (TR-dextran, 70 kDa, 10 mg/mL; **Supplemental Table 1**). The microinfusate spread from the microinstilled alveolus to airspaces of at least 20 alveoli, as evident by transient loss of optical discrimination between airspaces and alveolar walls. Return of optical discrimination within seconds of the microinstillation signaled that free microinfusate rapidly drained from alveoli toward airways to reestablish the air-filled alveolar lumens. In line with data published by our group and others (26, 30), subsequent confocal images showed fluorescence of the remaining TR-dextran as a juxta-epithelial layer that pooled at structural niches of alveoli, where septa converge (**Figure 1C**). Follow-up microinstillation of non-fluorescent vehicle solution into the TR-dextran-containing airspaces abolished the TR-dextran fluorescence (data not shown), indicating that TR-dextran was restricted to airspaces and not taken up by alveolar walls.

To define the effect of lung perfusion on alveolar liquid transport, we quantified time-dependent changes of TR-dextran fluorescence in alveolar airspaces of lungs that were perfused at PAP 15, 10, or 5 cm H_2_O and in lungs in which perfusion was halted (**Figure 1, D-F**). We focused our analysis on fluorescence of TR-dextran that pooled at alveolar corners (**Figure 1C**), where dextran volumes were larger and easier to visualize. We and others have shown previously that, provided alveolar barrier function is intact, time-dependent decrease of airspace dextran fluorescence intensity signals dextran dilution by the non-fluorescent alveolar wall liquid (AWL) secreted by the alveolar epithelium (26, 30). By contrast, time-dependent increase of airspace dextran fluorescence intensity signals airspace liquid absorption (26, 30). Before starting our experiments, we confirmed that change of TR-dextran fluorescence is an indicator of solution dilution and concentration, since calibration experiments in glass micropipettes showed TR-dextran fluorescence varies with dextran concentration in aqueous solution (**Supplemental Figure 2, A-B**).

In lungs perfused at PAP 15 cm H_2_O, we observed time-dependent loss of airspace TR-dextran fluorescence (**Figure 1, D-F**), suggesting the TR-dextran-containing solution was diluted by secreted AWL. TR-dextran fluorescence was unchanged by repeated exposure to confocal laser illumination (data not shown), ruling out the possibility that TR-dextran fluorescence loss was caused by photobleaching. Together, these findings confirm the published data (26, 30) and support the notion that the alveolar epithelium secretes AWL under baseline conditions.

Reports indicate that secreted AWL flows toward airways and does not accumulate in alveoli (30). In line with the published data (30), our findings show that widths of TR-dextran-containing liquid pools in alveoli did not change over time in lungs perfused at PAP 15 cm H_2_O (**Figure 1, G-H**), indicating there was no accumulation of airway liquid in alveoli to increase the volume of the alveolar surface liquid during the imaging period. Airspace diameter was also stable (**Figure 1, G-H**), indicating that our measurement of TR-dextran liquid width was not confounded by changes of alveolar morphology. These findings again confirm the published data (26, 30) and provide additional evidence that the alveolar epithelium secretes AWL in lungs perfused at PAP within the physiological range.

To evaluate the possibility that the observed TR-dextran fluorescence loss was caused by fluid leak from microvessels into airspaces, we defined alveolar barrier properties by adding fluorescein isothiocyanate (FITC)-tagged dextran (FITC-dextran, 20 kDa, 5 mg/mL) to the lung perfusate solution. We have shown previously that fluorophore-tagged dextran in the perfusate solution is retained in microvessels of uninfected alveoli, but it leaks into airspaces of lungs infected with *S. aureus* (SA), where SA-epithelial interactions cause alveolar barrier dysfunction (25, 26). We now affirmed that fluorescence of FITC-dextran in the perfusate solution was excluded from alveolar airspaces in uninfected lungs, but not in SA-infected lungs (**Figure 1, I-K**), indicating the alveolar barrier prevented airspace edema formation under baseline conditions. In addition, there was no widening of alveolar septa in uninfected lungs to suggest fluid accumulated in the alveolar interstitium (data not shown). We conclude that alveolar barrier function is intact in isolated, perfused lungs. Importantly, these data indicate that the airspace TR-dextran fluorescence loss we observed in lungs perfused at PAP 15 cm H_2_O (**Figure 1, D-F**) resulted from AWL secretion – not alveolar barrier leak.

Like alveoli of lungs perfused at PAP 15 cm H_2_O, alveoli of lungs perfused at PAP 10 cm H_2_O also showed time-dependent loss of airspace TR-dextran fluorescence (**Figure 1, D-F**), steady TR-dextran pool width (**Figure 1H**), steady airspace diameter (**Figure 1H**), and absence of FITC-dextran fluorescence in airspaces (**Figure 1K**). These findings were replicated in lungs of female mice (**Supplemental Figure 3, A-B**), indicating that mouse sex did not determine whether alveoli secreted AWL. Together, these findings show alveoli secrete AWL in lungs perfused at PAP that is near the physiological range. Notably, the rate of TR-dextran fluorescence loss in lungs perfused at PAP 10 cm H_2_O was not different from that of lungs perfused at PAP 15 cm H_2_O (**Figure 1, E-F**), suggesting AWL secretion is a regulated process for which ion transport across the alveolar epithelium is the rate-limiting step.

To test whether decreasing lung perfusion to PAP well below the physiological range alters alveolar liquid transport, we repeated the dextran experiments in lungs perfused at PAP 5 cm H_2_O and in lungs in which we halted lung perfusion. In both conditions, we observed time-dependent gain of airspace TR-dextran fluorescence (**Figure 1, D-F**), suggesting alveoli absorbed liquid from airspaces. In line with this possibility, TR-dextran pool width decreased over the imaging period while airspace diameter was steady (**Figure 1, G-H**). There was no leak of FITC-dextran from the perfusate solution into the airspaces in lungs perfused at PAP 5 cm H_2_O (**Figure 1K**) or in lungs in which perfusion was halted for 30 min, then reinitiated at PAP 10 cm H_2_O (data not shown). Taking these findings together, we conclude that halting lung perfusion or reducing it to PAP 5 cm H_2_O stimulated the alveolar epithelium to reverse liquid transport from homeostatic secretion to absorption.

To define the time course over which alveoli responded to lung hypoperfusion, we carried out an experiment in which we toggled lung perfusion between PAP 10 cm H_2_O and 5 cm H_2_O during a single imaging period (**Figure 2A**). Confocal images (**Figure 2, B-C** and **Supplemental Figure 4**) and plots (**Figure 2D**) show fluorescence and width of a representative pool of TR-dextran at an alveolar niche. Our findings show TR-dextran fluorescence decreased, but pool width was steady during the first imaging period at PAP 10 cm H_2_O (**Figure 2, C-D**), indicating alveoli secreted AWL. By contrast, within minutes of reducing PAP to 5 cm H_2_O, TR-dextran fluorescence increased while pool width decreased (**Figure 2, C-D**), indicating that reducing lung perfusion to PAP well below the physiological range rapidly stimulated airspace liquid absorption. Raising the perfusion back to PAP 10 cm H_2_O again caused TR-dextran fluorescence to decrease (**Figure 2, C-D**), indicating alveoli again secreted AWL. We observed TR-dextran pool width now increased (**Figure 2, C-D**), suggesting that raising the perfusion back to PAP 10 cm H_2_O stimulated AWL secretion that re-expanded the alveolar surface liquid volume. Reducing perfusion back to PAP 5 cm H_2_O caused TR-dextran fluorescence to increase and pool width to decrease (**Figure 2, C-D**), signaling alveoli absorbed airspace liquid. Taken together, these data show alveoli rapidly and reversibly switched between airspace liquid secretion and absorption in response to changes of lung perfusion. In addition, since both TR-dextran loss and gain repeatedly occurred in the same alveoli in the presence of continuous lung perfusion, these data rule out the possibility that TR-dextran loss in lungs perfused at or near physiological PAP resulted from alveolar barrier leak.

**Figure 2.**
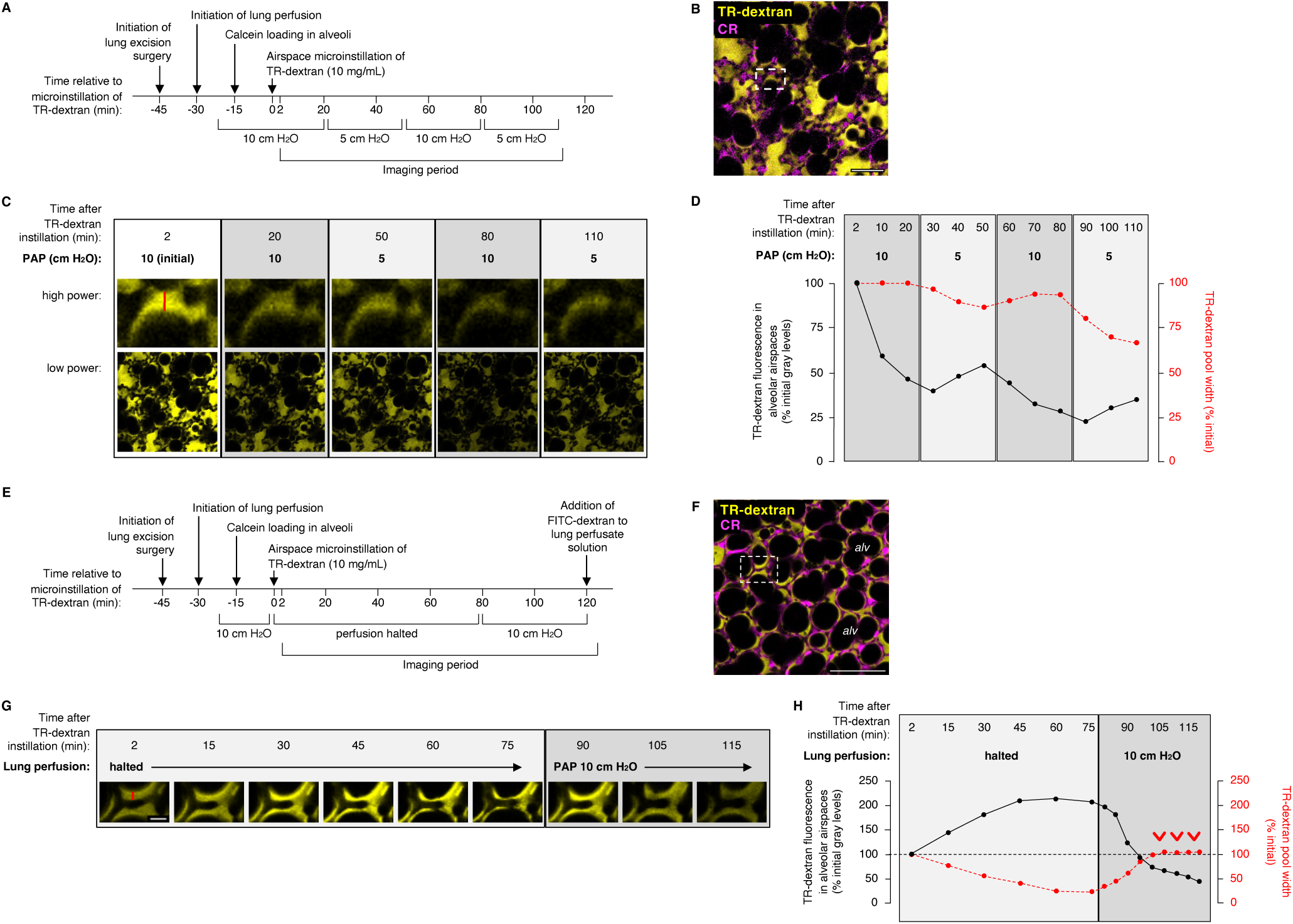
Alveoli respond rapidly to lung hypoperfusion. (**A-D**) Cartoon in <u>A</u> shows the design of experiments used to generate the confocal imaging data shown in B-D. The low power image in <u>B</u> shows a field of live alveoli (*magenta*) at 2 min after alveolar microinstillation of TR-dextran (*yellow*). The alveolar epithelium is delineated by fluorescence of calcein red-orange (*CR*), and the *dashed rectangle* indicates the location of the high power images and plots in C-D. The images and plots in <u>C-D</u> show change of TR-dextran fluorescence (*black plot*) and TR-dextran pool width (*red plot*), where fluorescence was quantified in the TR-dextran pool shown in the high power images (*top row*) and pool width was quantified at the location indicated by the *red line*. Note, TR-dextran fluorescence decreased during periods of lung perfusion at pulmonary artery pressure (*PAP*) 10 cm H2O, but increased during perfusion at PAP 5 cm H_2_O. The example images were replicated in 3 trials of alveolar TR-dextran microinstillation in imaging fields of at least 50 alveoli. Scale bar: 50 µm. **(E-H)** The cartoon in <u>E</u> shows the design of experiments used to generate the confocal imaging data shown in F-H. The low power image in <u>F</u> shows a field of live alveoli at 2 min after alveolar microinstillation of TR-dextran. The *dashed rectangle* indicates the location of the high power images and plots in G-H. The images and plots in <u>G-H</u> show change of TR-dextran fluorescence (*black plot*) and TR-dextran pool width (*red plot*), where fluorescence was quantified in the TR-dextran pool shown in the images and pool width was quantified at the location indicated by the *red line.* Note, after lung perfusion was restarted at PAP 10 cm H_2_O, TR-dextran pool width increased, then stabilized as pointed out by the *red arrowheads*. Scale bar: 100 (F) and 10 (G) µm.

The transient increase of TR-dextran meniscus width we observed during the second period of lung perfusion at PAP 10 cm H_2_O (**Figure 2, C-D**) raises the possibility that alveoli maintain a steady state airspace liquid height, as cultured alveolar epithelial cells do (31). To test this possibility, we carried out an experiment in which we halted, then restarted lung perfusion during a single imaging period (**Figure 2E**). Confocal images (**Figure 2, F-G**) and plots (**Figure 2H**) show fluorescence and width of a representative pool of TR-dextran at an alveolar niche. Our findings show TR-dextran fluorescence increased and pool width decreased during the period in which lung perfusion was halted (**Figure 2, G-H**), affirming that withholding lung perfusion stimulated alveolar liquid absorption. Restarting perfusion at PAP 10 cm H_2_O led to rapid decrease of TR-dextran fluorescence and increase of pool width (**Figure 2, G-H**), indicating that restoring perfusion to near physiological levels stimulated AWL secretion that expanded the alveolar surface liquid volume. Adding FITC-dextran to the lung perfusate at the conclusion of the experiment showed that withholding lung perfusion for 75 min did not lead to alveolar barrier leak, though barrier leak occurred in lungs in which we withheld lung perfusion for an additional 30 min (data not shown). We note that the TR-dextran pool width stabilized over time after perfusion was restored, and the final pool width was similar to the pool width at the time that lung perfusion was stopped (**Figure 2H**). These data suggest that, like cultured alveolar epithelial cells, the alveolar epithelium regulates AWL at a steady state liquid height or volume.

### Osmotic gradients near the physiological range do not determine liquid transport in alveoli

We next sought to define mechanisms of alveolar liquid secretion and absorption. We considered that trans-alveolar osmotic gradients between airspace and microvascular lumens might drive liquid transport in alveoli, based on published data (32–35). Our findings show the TR-dextran-containing airspace solution we used to generate the data shown in **Figures 1-2** was hyperosmolar to the lung perfusate solution (**Figure 3A** and **Supplemental Table 1**), raising the possibility that the trans-alveolar osmotic gradient present in our preparation, which is favorable to alveolar liquid secretion, was the driving force behind the TR-dextran fluorescence loss we observed in lungs perfused at PAP 15 and 10 cm H_2_O (**Figure 1, D-F**).

**Figure 3.**
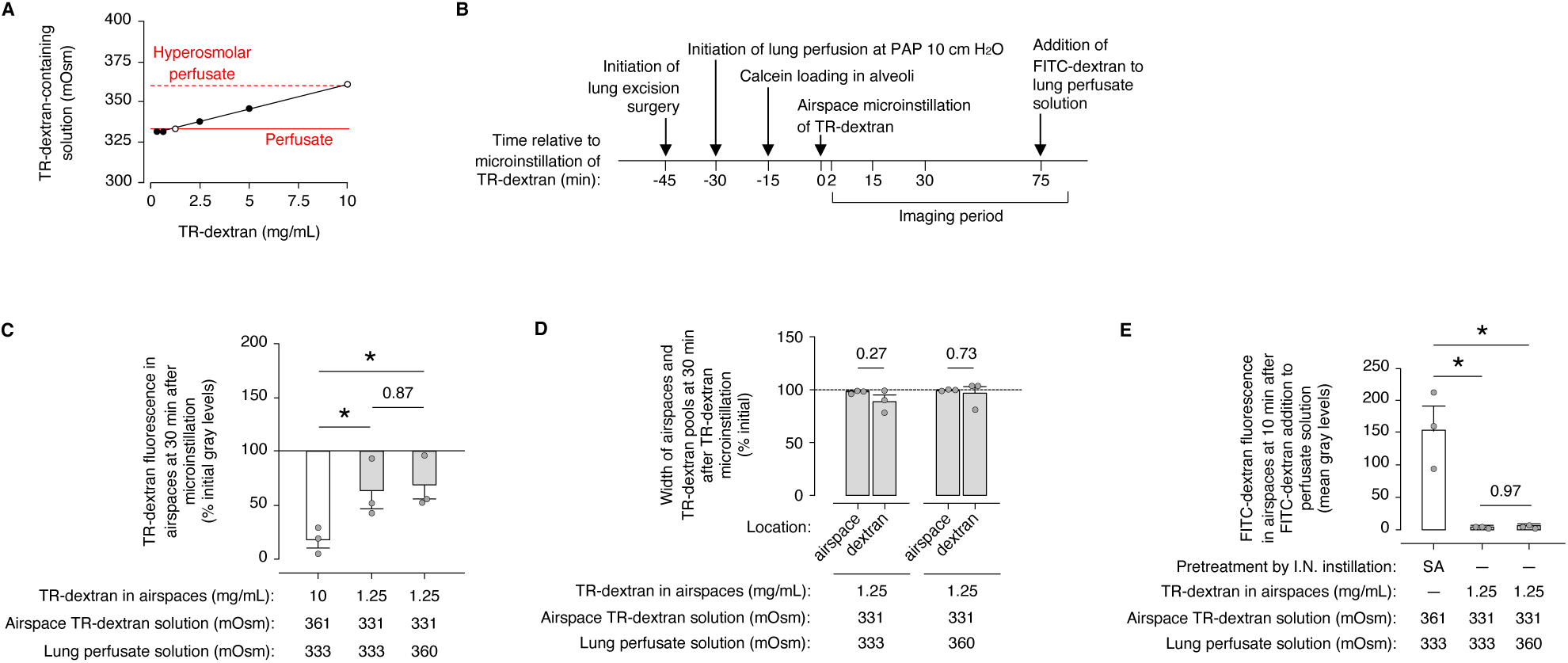
Trans-alveolar osmotic gradients that are near the physiological range do not determine the direction of liquid transport in alveoli. (**A-E**) The plot in <u>A</u> shows the effect of TR-dextran (70 kDa) concentration on solution osmolality. *White circles* indicate the TR-dextran-containing solutions that were microinstilled into alveolar airspaces to carry out the experiments shown in B-E. The *dashed and solid red lines* show osmolalities of the lung perfusate solutions, which did not contain TR-dextran, that were used in the same experiments shown in B-E. The cartoon in <u>B</u> shows the design of experiments used to generate the confocal imaging group data shown in C-E. Group data in <u>C-D</u> show change of alveolar TR-dextran fluorescence (C), TR-dextran pool width (D), and airspace width (D) at 30 min after microinstillation of the indicated TR-dextran solutions in live alveoli of lungs perfused with solutions of the indicated osmolalities. Group data in <u>E</u> show that FITC-dextran that we added to the lung perfusate solution leaked into alveolar airspaces only in lungs pretreated with intranasal (*I.N.*) instillation of *S. aureus* (SA). Data represented by the *white bars* in C and E are reproduced from Figures 1F and 1K, respectively. In C-E, *circles* indicate n and each represent 1 mouse in which mean change of fluorescence and width was quantified in imaging fields of at least 50 alveoli; *bars* show mean ± SEM; \**p* < 0.05 or as indicated by ANOVA with post hoc testing (C,E) and two-tailed *t* test (D).

To follow up, we quantified TR-dextran fluorescence changes in alveoli of lungs that we perfused at PAP 10 cm H_2_O but exposed to modified TR-dextran and perfusate solutions that were designed to abolish the original pro-secretion osmotic gradient, and, in separate experiments, reverse it to generate a pro-absorption gradient (**Figure 3, A-B** and **Supplemental Table 1**). The modified TR-dextran solution had the same components as the original TR-dextran solution, but the TR-dextran concentration was decreased (**Supplemental Table 1**). The modified perfusate solution was generated by adding sucrose to increase the solution osmolality to 360 mOsm/L, which is near the physiological range of serum osmolality in mice exposed to dehydration (**Supplemental Table 1**) (36, 37).

Our findings show airspace TR-dextran fluorescence decreased, but TR-dextran pool width and alveolar airspace width were steady in alveoli in which the airspace solution osmolality matched the lung perfusate osmolality (**Figure 3, C-D**). These findings show AWL secretion occurred in the absence of a pro-secretion osmotic gradient. We observed the same results in lungs in which the airspace solution osmolality was less than the lung perfusate osmolality (**Figure 3, C-D**), indicating that AWL secretion persisted in the presence of a pro-absorption gradient. FITC-dextran did not leak from microvessels into airspaces in any experiment (**Figure 3E**), indicating alveolar barrier function was intact. Separate experiments that showed airspace TR-dextran fluorescence decreased in alveoli of lungs in which TR-dextran was instilled by nebulization, prior to lung excision (**Supplemental Figure 5, A-C**), ruled out the possibility that the airspace TR-dextran fluorescence loss was caused by the alveolar micropuncture method. Together, these findings show alveoli secreted AWL in the presence of a trans-alveolar osmotic gradient that favored secretion, in the presence of a gradient that favored absorption, and in the absence of a gradient. We note that the TR-dextran fluorescence loss was less in the alveoli exposed to no gradient or to a pro-absorption gradient than it was in alveoli exposed to a pro-secretion gradient (**Figure 3C**), suggesting that an unfavorable osmotic gradient slowed AWL secretion. We conclude that the direction of liquid transport by the alveolar epithelium occurs independently of trans-alveolar osmotic gradients that are near the physiological range, though osmotic gradients may influence the AWL secretion rate. We interpret that AWL secretion is likely driven by active ion transport across the alveolar epithelium.

### Liquid absorption depends on ion transport proteins in the alveolar epithelium

We next aimed to define the ion transport proteins involved in alveolar liquid transport (**Figure 4, A-F**). Our general approach was to expose alveoli of isolated, perfused mouse lungs to drug inhibitors of ion transport proteins by either airspace instillation or addition to the lung perfusate solution (**Figure 4, A-B**). Our rationale was that drugs instilled by the airspace route would inhibit ion transport proteins primarily on the apical surface of the alveolar epithelium, while drugs instilled by the perfusate route would act primarily on the basolateral surface of the alveolar epithelium. After drug exposure, we microinstilled alveolar airspaces with TR-dextran, then added FITC-dextran to the lung perfusate solution (**Figure 4A**). There was no leak of FITC-dextran from microvessels into airspaces (data not shown).

**Figure 4.**
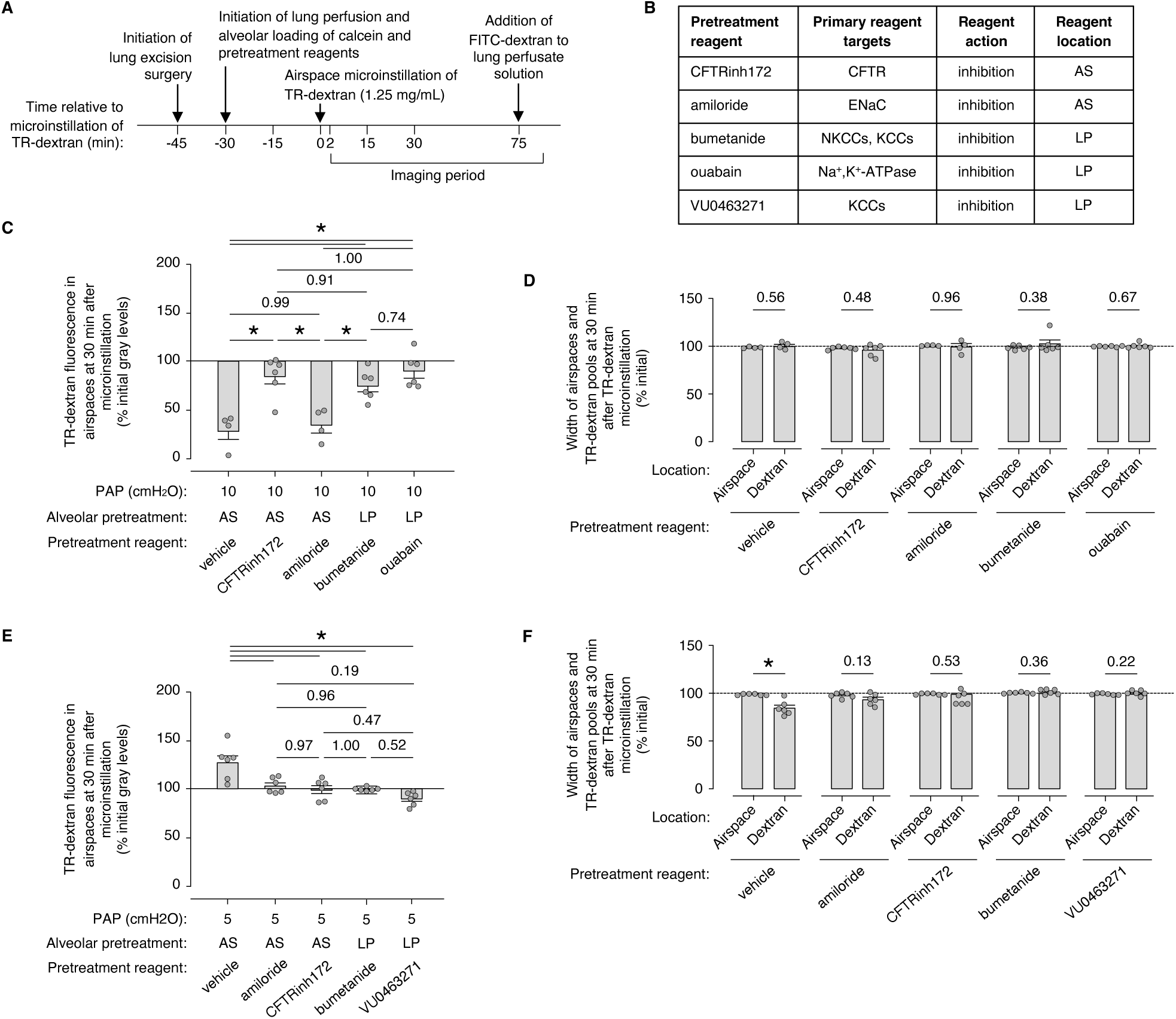
Ion transport protein mechanisms of alveolar liquid dynamics in live alveoli of perfused lungs. (**A-F**) Cartoon in <u>A</u> and reagent table in <u>B</u> show the design of experiments used to generate the group confocal imaging data shown in <u>C-F</u>. As indicated in A, we sequentially: 1. exposed the alveolar epithelium to solutions containing calcein and the pretreatment reagents indicated in B-F; 2. microinstilled the airspaces of the pretreated alveoli with TR-dextran (70 kDa); and 3. quantified change of TR-dextran fluorescence in alveolar airspaces at 30 min post-instillation. The alveolar epithelial exposures were by tracheal cannula instillation and confirmed by direct visualization of the solutions in alveoli. *AS*, airspace; *LP*, lung perfusate. Experiments were carried out in lungs perfused at PAP 10 cm H_2_O (C-D) or 5 cm H_2_O (E-F). In C-F, *circles* indicate n and each represent 1 trial of TR-dextran microinstillation in which mean fluorescence change was quantified in an imaging field of at least 50 alveoli; trials were carried out in lungs of at least 2 mice; *bars* show mean ± SEM; \**p* < 0.05 or as indicated by ANOVA with post-hoc Tukey testing (C,E) or two-tailed *t* test (D,F).

Experiments designed to define ion transport protein mechanisms of AWL secretion were carried out in lungs perfused at PAP 10 cm H_2_O (**Figure 4, A-D**). In alveoli in which we exposed the apical surface of the alveolar epithelium to vehicle solution, TR-dextran fluorescence decreased while TR-dextran pool width and airspace width were steady (**Figure 4, C-D**), as expected. Exposure of the apical surface of the alveolar epithelium to the CFTR inhibitor, CFTRinh172 blocked TR-dextran fluorescence loss but had no effect on TR-dextran pool width or airspace width (**Figure 4, C-D**). By contrast, apical exposure to the ENaC inhibitor, amiloride had no effect at all (**Figure 4, C-D**). Together, these findings show AWL secretion depends on CFTR in the alveolar epithelium but not ENaC, consistent with previously reported observations (26, 30).

Exposure of the basolateral surface of the alveolar epithelium to bumetanide, an inhibitor of NKCCs and CFTR, blocked TR-dextran fluorescence loss but did not affect TR-dextran pool width or airspace width (**Figure 4, C-D**). Since NKCC is located at the basal surface of the alveolar epithelium but CFTR is not, inhibition by bumetanide in the lung perfusate suggests NKCCs contribute to AWL secretion. We observed the same results in alveoli in which the basolateral surface of the alveolar epithelium was exposed to ouabain, an Na,K-ATPase inhibitor (**Figure 4, C-D**), indicating that AWL secretion is an active process driven by the alveolar epithelial Na,K-ATPase. Taking together all the findings generated in lungs perfused at PAP 10 cm H_2_O (**Figure 4, C-D**), we conclude that AWL secretion by the alveolar epithelium relies on CFTR-, NKCC-, and the Na,K-ATPase, but not ENaC. These conclusions are consistent with mechanisms of alveolar liquid secretion proposed previously by our group and others (22, 26, 30, 38).

Experiments designed to define ion transport protein mechanisms of alveolar liquid absorption were carried out in lungs perfused at PAP 5 cm H_2_O (**Figure 4, A-B and E-F**). In lungs in which we exposed the apical surface of the alveolar epithelium to vehicle solution, TR-dextran fluorescence increased while TR-dextran pool width decreased and airspace width was steady (**Figure 4, E-F**), as expected. Exposure of the apical surface of the alveolar epithelium to amiloride and CFTRinh172 each blocked the fluorescence increase and the pool width decrease (**Figure 4, E-F**). Taking these findings together, we conclude that alveolar liquid absorption in lungs perfused at PAP 5 cm H_2_O depends on ENaC and CFTR in the alveolar epithelium. These conclusions align with mechanisms of liquid absorption proposed by our group and others (22, 26, 38) and with the known capacity of CFTR to mediate both liquid secretion and absorption.

Exposure of the basolateral surface of the alveolar epithelium to bumetanide in the perfusate solution also blocked the TR-dextran fluorescence increase and the pool width decrease (**Figure 4, E-F**). We found these findings surprising, since they seemed to implicate NKCCs – ion transport proteins known for Na^+^ and Cl^-^ *import* into the alveolar epithelium – in a liquid absorption mechanism that should require Na^+^ and Cl^-^ *export* to the interstitium. We considered the alternative possibility that the bumetanide effect resulted from bumetanide-induced inhibition of K^+^-Cl^-^-cotransporters (KCCs), which are K^+^ and Cl^-^ exporters that are expressed by the alveolar epithelium and generally locate to epithelial cell basolateral surfaces (39, 40). To test whether KCCs contribute to alveolar liquid absorption in hypoperfused lungs, we exposed the basolateral surface of the alveolar epithelium to VU0463271, a drug that inhibits KCCs but not NKCCs (41, 42). Our findings show VU0463271 blocked the TR-dextran fluorescence increase and the pool width decrease (**Figure 4, E-F**), indicating that KCCs are central to the mechanism of hypoperfusion-induced alveolar liquid absorption. Some loss of airspace TR-dextran fluorescence in alveoli of VU0463271-exposed lungs (**Figure 4E**) raises the possibility that VU0463271 not only inhibited alveolar liquid absorption, but also restored AWL secretion by mechanisms that remain unclear. We conclude that alveolar liquid absorption in hypoperfused lungs depends on alveolar epithelial KCCs, CFTR, and ENaC.

### Hypoperfusion causes changes of alveolar structure

We next sought to understand how lung hypoperfusion stimulated a switch of alveolar liquid transport from secretion to absorption. Reports indicate that static reductions of lung perfusion pressure decrease alveolar microvascular lumen diameters (19), indicating that lung hypoperfusion alters alveolar morphology. We considered that hypoperfusion-induced changes of alveolar morphology might inhibit NKCC-mediated AWL secretion and stimulate KCC-mediated alveolar liquid absorption through alveolar epithelial stretch. Our reasoning derives from two observations: first, alveolar epithelial cell stretch alters epithelial ion transport in vitro (43); and second, cell stretch inhibits NKCCs and activates KCCs in cultured cell models of cell volume regulation (44–46).

To determine how lung hypoperfusion alters alveolar morphology, we measured microvessel and airspace lumen diameters in alveoli exposed to a step-wise decrease, then increase of lung perfusion in the same imaging period (**Figure 5A**). Specifically, we initiated lung perfusion at PAP 15 cm H_2_O, decreased it to the extreme point at which perfusion was withheld, then restarted it and increased it back to PAP 15 cm H_2_O (**Figure 5A**). Confocal images show that each 5 cm H_2_O decrease of PAP contracted alveolar microvascular lumen diameters, while each 5 cm H_2_O increase of PAP expanded them (**Figure 5, B-C** and **Supplemental Figure 6**). These findings confirm reports indicating that reduction of lung perfusion decreases microvessel lumen size (19). In addition, these data add the observation that restoration of lung perfusion rapidly reestablishes microvessel lumens. Interestingly, the final measures of microvascular lumen diameters in the post-reperfusion phase exceeded the initial values (**Figure 5, B-C** and **Supplemental Figure 6**), suggesting that the microvascular endothelium had a vasodilatory response to the short, but profound period of lung hypoperfusion.

**Figure 5.**
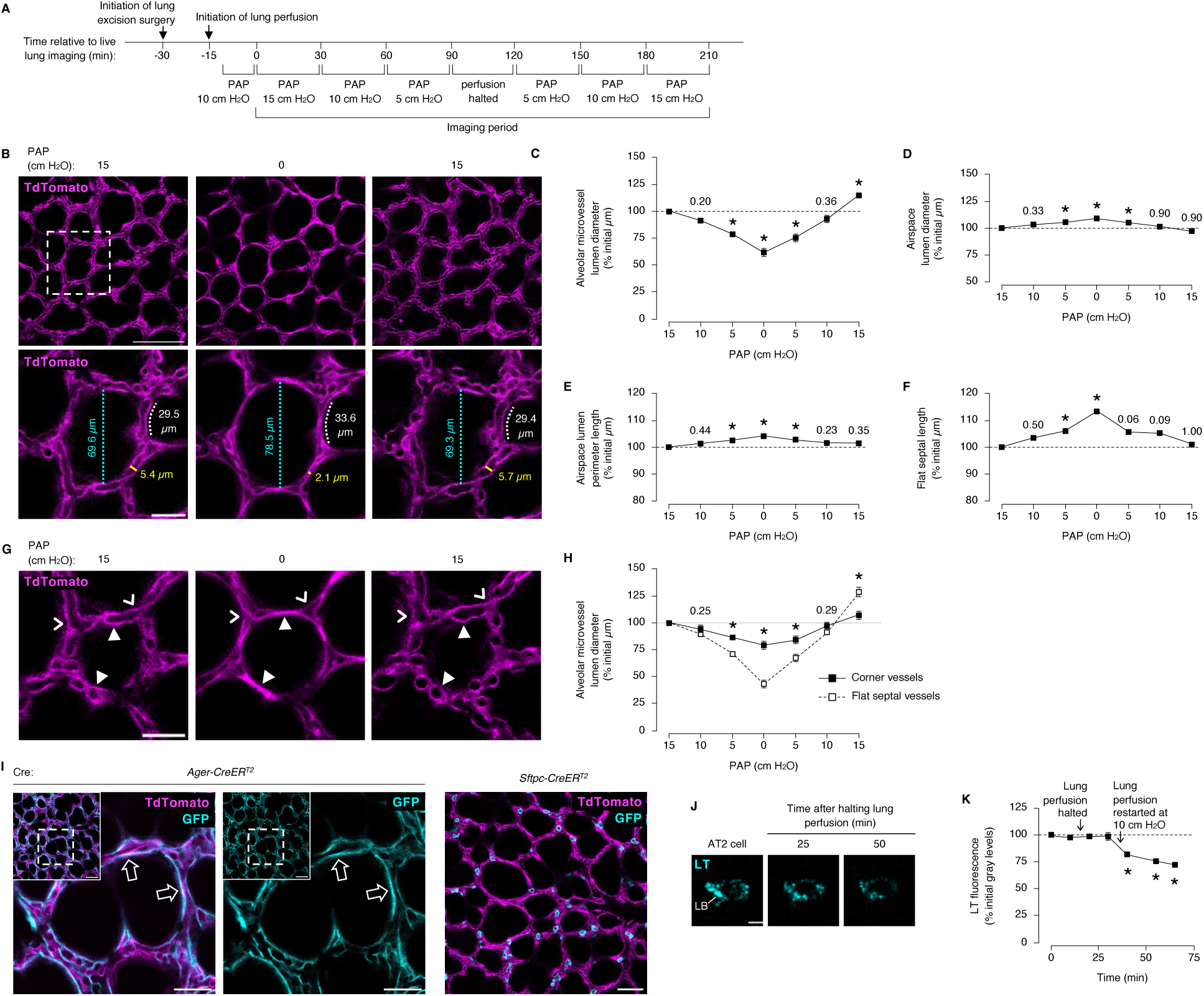
Lung hypoperfusion causes alveolar epithelial stretch. (**A-H**) Cartoon in <u>A</u> shows the design of experiments used to generate the confocal imaging data in B-H. Images were generated in live, perfused lungs of a *ROSA^mT/mG^*-expressing mouse in which the alveolar epithelium and endothelium are marked by TdTomato fluorescence (*magenta*). In <u>B</u>, the *dashed rectangle* shows the location of the high power images in the *bottom row*. Images and group data in <u>B-F</u> show changes of alveolar microvascular lumen diameter (C, *yellow lines and text* in B), airspace lumen diameter (D, *cyan lines and text* in B), lumen perimeter (E), and septal length (F, *white lines and text* in B) in response to changes of pulmonary artery pressure (*PAP*). In C-F, *squares* represent mean ± SEM of measurements in 4 imaging fields of at least 50 alveoli; \**p* < 0.05 or as indicated by ANOVA with post hoc Tukey testing versus the *first square.* Images and group data in <u>G-H</u> show changes of microvascular diameter at flat segments of alveoli (*closed arrowheads*) and alveolar corners (*open arrowheads*). In H, *squares* are as in C-F, but \**p* < 0.05 or as indicated versus flat septal vessels by two-tailed *t* test. Scale bars: 100 (B, *top row*), 30 (B, *bottom row*), and 25 (G) µm. **(I)** High (*left, center*) and low (*insets, right*) power images show alveoli of live, perfused lungs of *ROSA^mT/mG^*– and *Ager-CreER^T2^*or *Sftpc-CreER^T2^*-expressing transgenic mice. *GFP* marks AT1 or AT2 cells, respectively. Note, AT1 cells line flat septa (*open arrows*), while AT2 cells occupy corners. Images replicated in 3 mice per group. Scale bars: 50 (*inset, right*) and 25 (*left, center*) µm. **(J-K)** Images and plot show the effect of halting lung perfusion for 15 min on fluorescence of AT2 cell lamellar bodies (*LB*) loaded with lysotracker dye (*LT*). See text. *Squares:* mean ± SEM of 3 trials of LT microinstillation in which mean fluorescence was quantified in imaging fields of at least 20 AT2 cells. Images replicated in 2 mice. \**p* < 0.05 versus first value by ANOVA with post hoc Tukey testing. Scale bar: 5 µm.

Alveolar microvessels were not the only alveolar structure that showed a morphological response to lung hypoperfusion. Each 5 cm H_2_O decrease of PAP increased alveolar airspace lumen diameters, while each 5 cm H_2_O increase of PAP reduced them toward their original sizes (**Figure 5, B and D** and **Supplemental Figure 6**). We observed the same pattern when we quantified airspace lumen perimeters and alveolar septal lengths (**Figure 5, B and E-F** and **Supplemental Figure 6**).

Additional analysis of the confocal images shows that the most marked changes of alveolar morphology occurred at flat regions of alveolar septa, where microvessel lumen diameters decreased by more than half (**Figure 5, G-H**). Microvessels at alveolar corners were relatively protected (**Figure 5, G-H**). These findings align with published data (20) and suggest alveolar cells located at flat septa might be subjected to greater degrees of hypoperfusion-induced changes of alveolar structure. Follow-up imaging of live lungs of transgenic reporter mice, in which GFP expression marks alveolar type 1 (AT1) and type 2 (AT2) cells, showed that AT1 cells line flat alveolar septa while AT2 cells occupy alveolar corners (**Figure 5I**). Although the locations of AT1 cells at flat septa and AT2 cells at corners is an expected finding, these data raise the possibility that AT1 cells are major sites of hypoperfusion-induced morphological changes in alveoli.

### Hypoperfusion causes alveolar stretch that stimulates liquid absorption

Since reports indicate ion transport protein responses occur in cultured alveolar epithelial cells after a surface area increase of only 10-12% (43, 47) – a value similar to the septal length increase we identified after halting perfusion (**Figure 5F**) – we considered lung hypoperfusion might cause alveolar epithelial stretch that alters ion and liquid transport in alveoli. Alveolar epithelial stretch can be identified in real time in intact, perfused lungs using lysotracker dye, since loss of lysotracker fluorescence in AT2 cells signals stretch-induced surfactant release (25, 48, 49). Live lung imaging showed AT2 cell lysotracker fluorescence was steady in lungs perfused at PAP 10 cm H_2_O, but it rapidly decreased after perfusion was halted for 15 min, then restarted at PAP 10 cm H_2_O (**Figure 5, J-K**). These findings show lung hypoperfusion stimulated AT2 cell surfactant release. We conclude that lung hypoperfusion caused alveolar epithelial stretch.

Interestingly, lysotracker fluorescence decreased after lung perfusion was restarted (**Figure 5K**), suggesting that while alveolar stretch stimulated the surfactant secretion response, surfactant release occurred only after the stretch stimulus resolved. The delay of surfactant release could have been caused by factors related to hypoperfusion, such as loss of alveolar bioenergetics due to loss of perfusate glucose delivery to alveoli. Alternatively, the delay could have resulted from the prolonged stretch stimulus. To differentiate between these possibilities, we quantified AT2 cell lysotracker fluorescence in alveoli of lungs we subjected to a prolonged period of hyperinflation. Live lung imaging showed AT2 cell lysotracker fluorescence was steady in lungs inflated to airway pressure 6 cm H_2_O, remained steady during a 15 min hyperinflation to airway pressure 12 cm H_2_O, then decreased after airway pressure was returned to 6 cm H_2_O (**Supplemental Figure 7**). The pattern of lysotracker fluorescence change we identified in lungs subjected to prolonged hyperinflation mirrors the pattern we observed in hypoperfused lungs. We conclude that while the surfactant secretion response is initiated by hyperinflation– or hypoperfusion-induced alveolar stretch, surfactant release is delayed until the hyperinflation is resolved or a physiological level of perfusion is restored.

Finally, we considered that alveolar epithelial stretch was the stimulus that inhibited AWL secretion and initiated alveolar liquid absorption in hypoperfused lungs. To test the hypothesis that alveolar stretch causes alveolar liquid absorption, we induced alveolar epithelial stretch by hyperinflation to airway pressure 12 and 16 cm H_2_O in lungs perfused at PAP 10 cm H_2_O (**Figure 6, A-B**). Confocal images show lung hyperinflation caused AT2 cell surfactant secretion (**Figure 6A**) and expanded alveolar airspace lumen diameters and septal lengths (**Figure 6, B-E**). These findings are consistent with findings reported by others (50) and show that lung hyperinflation causes alveolar epithelial stretch. Microvessel lumen diameters did not change during the hyperinflation period (data not shown). In lungs hyperinflated to airway pressure 12 cm H_2_O, airspace TR-dextran fluorescence increased, TR-dextran pool width decreased, and airspace width was steady (**Figure 6, F-G**), indicating that hyperinflation stimulated liquid absorption from airspaces. Taken together, these findings show hyperinflation-induced epithelial stretch in alveoli blocked AWL secretion and stimulated alveolar liquid absorption. Since lung hypoperfusion caused alveolar epithelial stretch (**Figure 5, J-K**), we propose that stretch of the alveolar epithelium was the stimulus that initiated a reversal of alveolar liquid transport from homeostatic secretion to absorption in hypoperfused lungs (**Figure 7**).

**Figure 6.**
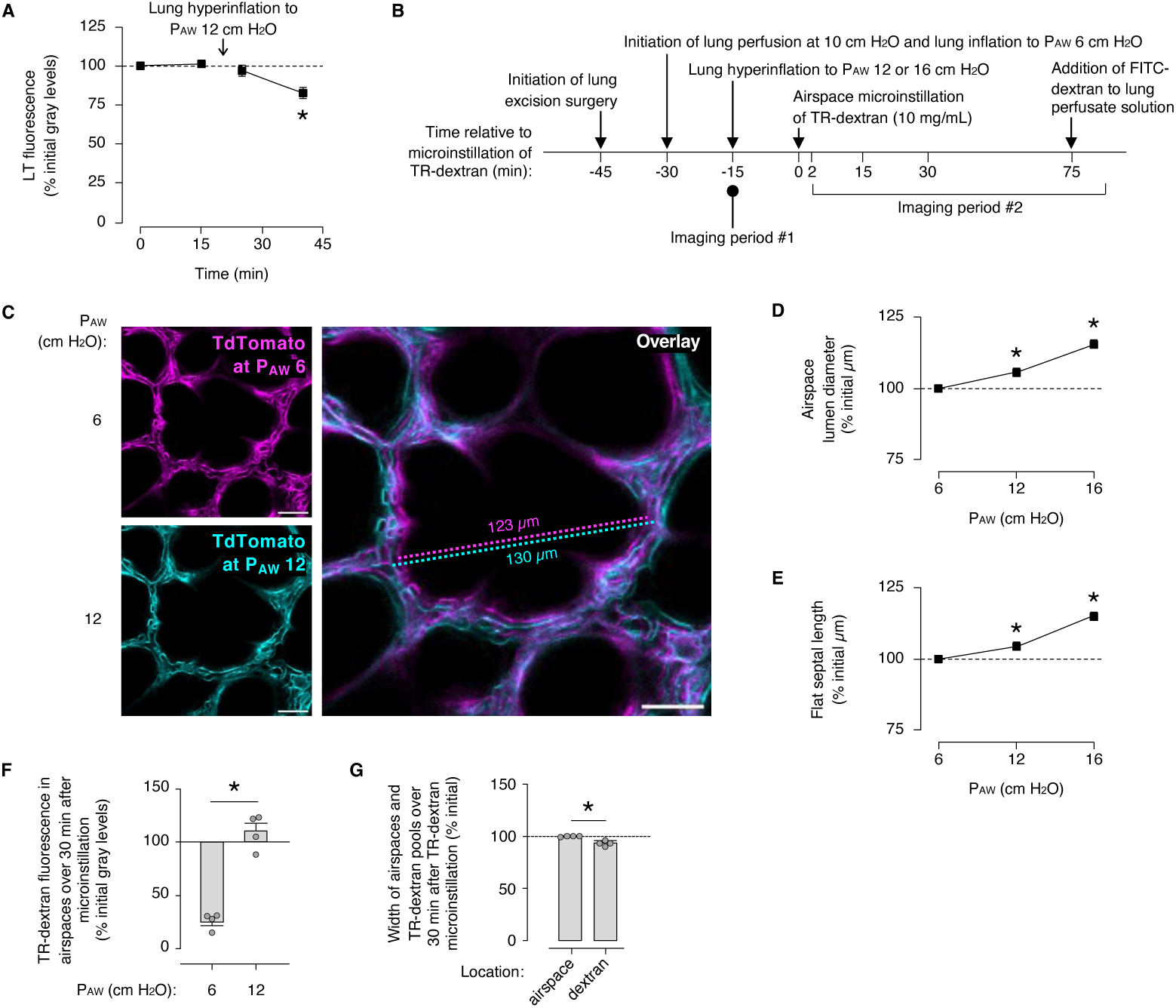
Alveolar epithelial stretch stimulates liquid absorption in alveoli. **(A)** Group confocal imaging data show the effect of a 15-second lung hyperinflation to airway pressure (P_AW_) 12 cm H_2_O on fluorescence of AT2 cell lamellar bodies loaded with lysotracker dye (*LT*). *Squares:* mean ± SEM of 4 trials of LT microinstillation in which mean fluorescence was quantified in imaging fields of at least 8 AT2 cells. \**p* < 0.05 versus *first square* by ANOVA with post hoc Tukey testing. **(B-G)** Cartoon in <u>B</u> shows the design of experiments used to generate the imaging data shown in C-G. Images were generated in live, perfused lungs of a *ROSA^mT/mG^*-expressing mouse in which TdTomato fluorescence marks the alveolar epithelium. Images and group data in <u>C-E</u> show hyperinflation-induced change of alveolar morphology. As indicated in B, we sequentially: 1. inflated the lungs to P_AW_ 6 cm H_2_O; 2. generated confocal images; 3. increased the P_AW_ to 12 or 16 cm H_2_O; and 4. generated confocal images again. Images of an example alveolus at P_AW_ 6 and 12 cm H_2_O were superimposed to generate the *overlay* in C*. Magenta pseudocolor, line, and text* show the diameter of the example airspace at P_AW_ 6 cm H2O, while *cyan pseudocolor, line, and text* show the diameter of the same airspace at P_AW_ 12 cm H_2_O. Images at P_AW_ 16 cm H_2_O are not shown. In D-E, *squares* represent mean ± SEM of measurements in 4 imaging fields of at least 50 alveoli; \**p* < 0.05 by ANOVA with post hoc Fisher testing versus *first square.* To generate the group data shown in <u>F-G</u>, we microinstilled TR-dextran (70 kDa) in alveolar airspaces of lungs held at P_AW_ 6 cm H_2_O (*left bar* in F) or 12 cm H_2_O (*right bar* in F, G). In F-G, *circles* indicate n and each represent 1 trial of TR-dextran microinstillation in which mean fluorescence change was quantified in an imaging field of at least 50 alveoli; trials were carried out in lungs of at least 2 mice; *bars* show mean ± SEM; \**p* < 0.05 by two-tailed *t* test.

**Figure 7.**
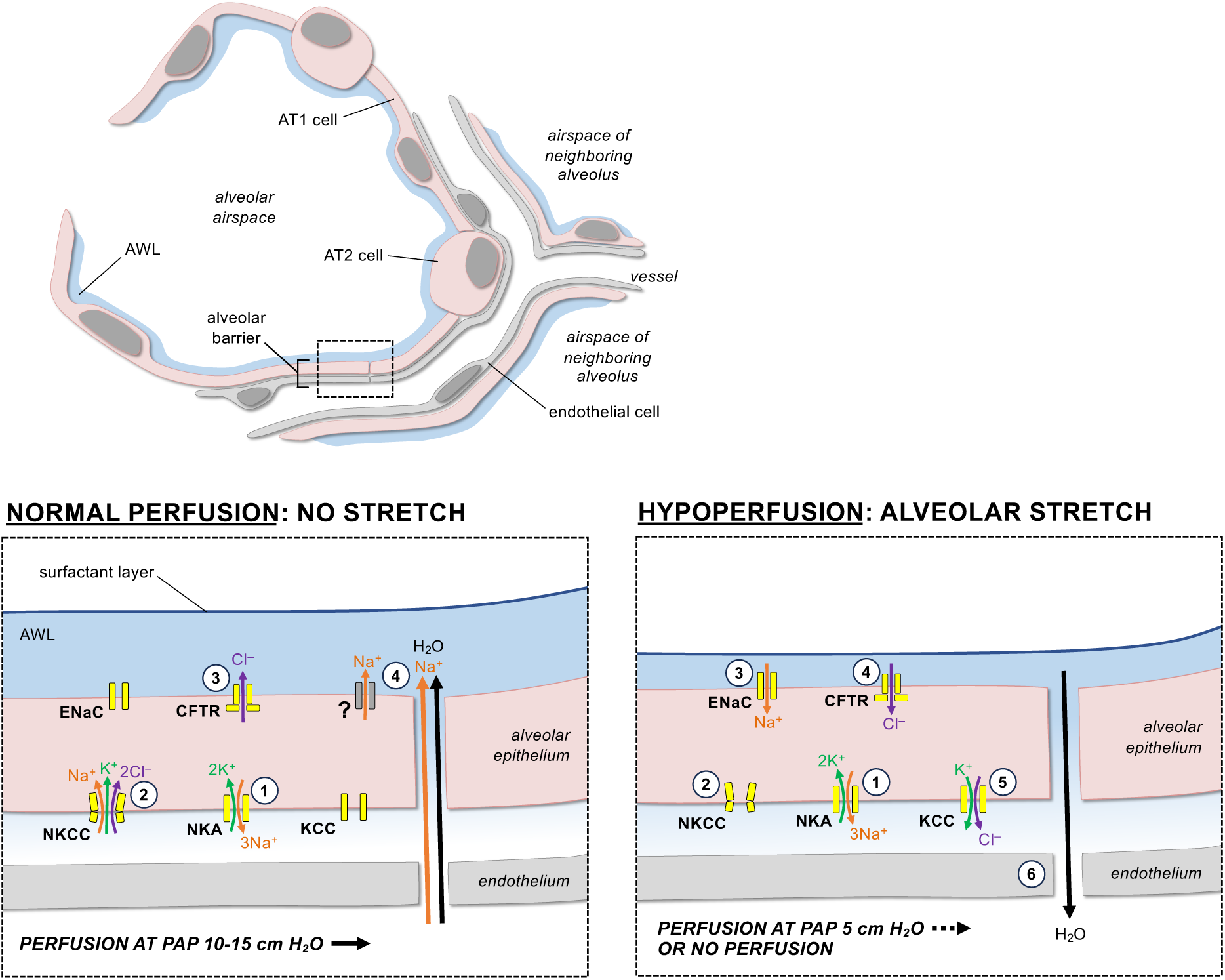
Proposed mechanism of hypoperfusion-induced liquid absorption in alveoli. Cartoons of an alveolus (*top*) and an alveolar wall (*bottom panels*; location shown by *dashed rectangle*) illustrate the proposed mechanisms in lungs perfused under conditions of normal perfusion or hypoperfusion as indicated. **<u>Left panel</u>:** The alveolar epithelium secretes alveolar wall liquid (*AWL*) in lungs perfused at physiological or near-physiological pulmonary artery pressures of 15 or 10 cm H_2_O, respectively. (**1**) Constitutive activity of the basolateral Na^+^/K^+^-ATPase (*NKA*) in the alveolar epithelium transports Na^+^ out of the cytosol and into the interstitial space, establishing a Na^+^ gradient across the basolateral plasma membrane. (**2**) The Na^+^ gradient drives epithelial import of Na^+^ and Cl^-^ through the basolateral Na^+^-K^+^-Cl^-^ cotransporter (*NKCC*). (**3**) Cl^-^ import increases intracellular Cl^-^ and establishes a Cl^-^ gradient that drives Cl^-^ export through apical CFTR and into airspace lumens. (**4**) Na^+^ and water follow by transcellular or paracellular pathways to generate AWL. **<u>Right panel</u>:** The alveolar epithelium absorbs airspace liquid in lungs exposed to hypoperfusion caused by either halting lung perfusion or reducing it to PAP 5 cm H_2_O, a level well below the physiological range. In alveoli of hypoperfused lungs, <u>shrinkage</u> <u>of microvascular lumens leads to expansion of airspace lumens that causes alveolar epithelial stretch</u>, which may inhibit NKCC-mediated ion transport and activate KCC-mediated ion transport. We propose: (**1**) constitutive Na^+^/K^+^-ATPase activity continues to establish a Na^+^ gradient across the basolateral plasma membrane; (**2**) NKCC-mediated Na^+^ import decreases, generating an intracellular Na^+^ gradient that (**3**) stimulates ENaC-mediated Na^+^ import; (**4**) Cl^-^ follows the airspace-to-interstitium path of Na^+^ via Cl^-^ import by apical CFTR and (**5**) Cl^-^ export by basolateral KCCs; and (**6**) water follows by paracellular pathways, reducing the alveolar lining liquid volume. Data supporting *left panel* steps 1-4 are provided in Figures 1-4; data supporting the notion that hypoperfusion causes alveolar epithelial stretch are provided in Figures 5-6; and data supporting *right panel* steps 3-6 are provided in Figures 1-4. Literature supporting *right panel* step 2 is provided in the Discussion.

## DISCUSSION

Our findings provide the first direct evidence, of which we are aware, that alveoli respond to lung hypoperfusion by reversing epithelial liquid transport from homeostatic secretion to absorption. Thus, while lung perfusion at or near physiological levels led to CFTR-, NKCC– and Na,K-ATPase-dependent AWL secretion by the alveolar epithelium, halting or reducing lung perfusion to a level well below the physiological range rapidly stimulated ENaC-, CFTR-, and KCC-dependent alveolar liquid absorption. Lung hypoperfusion associated with contraction of alveolar microvessel lumens and expansion of airspace lumens that caused alveolar epithelial stretch. Since stretch of the alveolar epithelium caused alveolar liquid absorption, we propose alveolar epithelial stretch was the stimulus that drove the switch to alveolar liquid absorption in hypoperfused lungs, perhaps by modulating NKCC and KCC activity. Notably, liquid and surfactant secretion remained coupled in alveoli of hypoperfused lungs. These findings inform the understanding of alveolar responses to systemic and local hypoperfusion. In addition, they raise the possibility that dysregulation of alveolar liquid transport and surfactant secretion have roles in the pathogenesis of lung diseases that are characterized by acute microvascular hypoperfusion, such as acute chest syndrome, severe viral lung infection, ventilator-induced lung injury, and acute respiratory distress syndrome.

Our findings reveal that lung alveoli, like other tissues, respond to hypoperfusion by absorbing liquid. Our discovery that lung hypoperfusion stimulated a reversal of alveolar liquid transport from secretion to absorption is novel, but it is well-supported by reports of similar findings in extrapulmonary tissues. For example, blood loss-induced hypotension in cats stimulates a switch from liquid secretion to absorption in the gut and skin (51). Low diastolic blood pressure stimulates liquid absorption in perfused rat intestine (52). Acute decrease of tissue blood flow in humans stimulates liquid absorption in forearm muscle and jejunum (53, 54). Why the lung, gut, skin, and muscle respond to hypoperfusion by absorbing liquid is not clear. One possibility is that absorption is a compensatory response that seeks to normalize tissue perfusion by augmenting intravascular liquid volume. Since the liquid volume in alveolar airspaces is less than 40 mL (55), alveolar liquid absorption alone is unlikely to restore tissue perfusion, but it might contribute to a global absorptive response to hypovolemia. Related questions remain. For instance, whether alveoli continue to absorb liquid in the face of prolonged hypoperfusion is not yet clear. Also, the extent to which the mechanisms that mediate hypoperfusion-induced liquid absorption are the same mechanisms that mediate reversals of alveolar liquid transport in response to other stimuli, such as influenza lung infection and hydrostatic stress (22, 26), is not known. Although future research might provide insights into these and other related questions, the findings included in this report provide strong evidence that lung alveoli respond to hypoperfusion with liquid absorption.

An important aspect of our data is the idea that alveolar epithelial stretch underpins liquid absorption mechanisms in hypoperfused lungs. A mechanistic role for alveolar stretch is supported by two lines of evidence. First, lung hypoperfusion increased alveolar diameters, perimeters, and septal lengths and stimulated surfactant secretion upon restoration of perfusion to physiological levels, suggesting that lung hypoperfusion caused sufficient stretch of the alveolar epithelium to initiate a physiological stretch response. Second, alveolar stretch by a separate mechanism – that is, lung hyperinflation – had the same effect as lung hypoperfusion, in that it increased alveolar diameters and septal lengths and initiated alveolar liquid absorption. Together, these findings provide evidence that lung hypoperfusion caused alveolar epithelial stretch that stimulated liquid absorption. Other factors might have contributed to the directional change of alveolar liquid transport that occurred in hypoperfused lungs. For example, hypoperfusion-induced reductions of microvascular flow and lumen diameter probably reduced shear stress and radial outward force on the microvascular endothelium. If changes of shear stress and radial force caused an endothelial response, endothelial-epithelial communication via ROS (56) or other diffusible factors could have contributed to epithelial liquid absorption. Microenvironmental factors like hypoxia might have also played a role in lungs in which perfusion was halted. Going forward, future studies might clarify the extent to which microenvironmental factors and the endothelium participate in mechanisms of hypoperfusion-induced liquid absorption in alveoli. Whether or not the endothelium has a role, we point out that the alveolar epithelium must be central to the liquid absorption mechanism, since the fluid– and solute-impermeable nature of the alveolar epithelial barrier (57, 58) isolates the endothelium from the epithelial site of airspace liquid absorption.

Our discovery that hypoperfusion-induced alveolar liquid absorption was KCC-dependent provides an additional layer of support for alveolar epithelial stretch as a critical absorption-inducing stimulus. Alveolar epithelial cells express multiple KCC isoforms including KCC1 and KCC4, and KCCs in the basolateral membranes of alveolar epithelial cells have been proposed previously to form a pathway, along with apical membrane CFTR, for apical-to-basolateral Cl^-^ flux (38, 39). Our findings show that KCCs were critical to alveolar liquid absorption in hypoperfused lungs, while NKCCs were required for AWL secretion in lungs perfused at or near physiological levels. KCCs and NKCCs are not mechanosensitive proteins, but they have reciprocal roles in cellular responses to mechanical stimuli. For example, cell swelling leads to NKCC and KCC dephosphorylation that inactivates NKCCs but activates KCCs, leading to decreased cellular Na^+^ import, increased K^+^ and Cl^-^ export, and compensatory reduction of intracellular fluid cell volume (44–46). Cell shrinkage has the opposite effect (44–46). Bringing our findings together with the published data, we propose that in lungs perfused at or near physiological levels, NKCCs facilitate ion transit in the alveolar epithelium that leads to AWL secretion (**Figure 7**). However, in lungs in which perfusion is halted or well below the physiological range, stretch of the alveolar epithelium in response to reduction of microvascular lumen diameter might cause NKCC inhibition and KCC activation, thereby stimulating a change of epithelial ion transport that favors airspace liquid absorption (**Figure 7**). In summary, these findings identify that KCCs were critical to hypoperfusion-induced alveolar liquid absorption, and they call new attention to KCCs as potential mediators of lung health and disease.

In our studies, perfusion-related changes of alveolar structure affected the alveolar epithelium in a spatially heterogeneous manner that might lead to epithelial stretch responses that are cell type-specific. In this regard we note two points. First, we identified that microvessel lumens at flat regions of alveoli contracted to a greater extent than microvessel lumens at alveolar corners, suggesting flat septa are more vulnerable to the stretch effects of lung hypoperfusion than alveolar corners, which are relatively protected. These findings reproduce published data that show lung perfusion determines alveolar microvascular lumen diameter (19, 20) and that microvessels at alveolar corners retain patency under conditions of hypoperfusion (20, 59), likely due to the structural system of lung connective tissue fibers that exerts tension on corner microvessels. Second, we showed that flat regions of alveoli are lined primarily by AT1 cells, raising the possibility that AT1 cells preferentially sense hypoperfusion-induced alveolar epithelial stretch and might initiate the switch from AWL secretion to airspace liquid absorption. Findings published by others support a potential role for AT1 cells in alveolar responses to stretch, since reports indicate that flat septa are sites of linear distention in alveoli of hyperinflated lungs (50), and that hyperinflation stimulates Ca^2+^ signaling that initiates in AT1 cells (49). A role for AT1 cells is further supported by data demonstrating that AT1 cells express CFTR, ENaC, and Na^+^/K^+^-ATPase (60–62) and may contribute to AWL secretion (63). Future investigations that compare microanatomical alveolar regions and alveolar epithelial cell types might shed light on how flat septa and alveolar corners differ in terms of their ion transport protein compositions, stretch sensitivity, and physiological responses to lung hypoperfusion. In addition, it remains an open question whether alveoli located in specific lung regions – for example, Zone 1 as identified by West and colleagues, where alveolar pressure exceeds pulmonary arterial pressure (64) – are more susceptible to hypoperfusion-induced changes of alveolar morphometry, as suggested by Ciurea and Gil (20). Our findings support the possibility that alveoli in Zone 1 conditions could be more exposed to hypoperfusion-induced alveolar stretch and, thus, respond with alveolar liquid absorption.

Since we identified that alveoli of lungs perfused at or near physiological PAP secrete AWL, our data strengthen the evidence that liquid secretion by the alveolar epithelium generates the alveolar lining layer at homeostasis. The extent to which alveoli secrete or absorb liquid under homeostatic conditions is controversial (65). The notion that the alveolar epithelium secretes liquid under baseline conditions is supported by real-time imaging of live alveoli of intact, perfused lungs, which show fluorescence of fluorophore-tagged dextran in alveolar airspaces is lost over time in the absence of an extreme, pro-absorption osmotic gradient (26, 30, 32–35). Our new findings add that airspace TR-dextran fluorescence was lost in alveoli of lungs perfused at or near physiological levels, irrespective of whether a physiologic-range osmotic gradient favored secretion, absorption, or no liquid movement. Our observation that airspace dextran fluorescence decreased, then increased in the same alveoli in response to perfusion changes in lungs that were perfused continuously align with reports that show alveolar barrier function is intact in isolated, perfused lungs (66, 67) and provide strong evidence against the possibility that dextran fluorescence loss in our experiments was caused by alveolar barrier leak. The CFTR-, NKCC-, and Na^+^/K^+^-ATPase-dependent mechanisms of alveolar liquid secretion we identified align with mechanisms put forth by others in the lung (22, 30, 38) and non-lung tissues (68). We conclude that our findings support previous suggestions that the alveolar epithelium secretes liquid at homeostasis (30, 69, 70), though it remains unclear where the liquid flows and to what extent it might be absorbed in the small airways. Taking our new findings together with the published data, we conclude that the alveolar epithelium secretes liquid under baseline conditions to generate the aqueous portion of the alveolar lining layer.

Our observation that surfactant release was blocked during the period of lung hypoperfusion suggests that coupling between liquid and surfactant secretion was maintained in alveoli of hypoperfused lungs. Since alveolar stretch is a major stimulus for AT2 cell surfactant secretion, it seems perhaps puzzling that hypoperfusion blocked surfactant secretion. The current understanding of surfactant secretion mechanisms is that transient, hyperinflation-induced stretch of the alveolar epithelium is sensed by AT1 cells, which respond by initiating Ca^2+^ signals that are conducted to AT2 cells (49, 71). Subsequent increases of cytosolic Ca^2+^ in AT2 cells leads to cytoskeletal changes that facilitate lamellar body fusion with AT2 cell plasma membranes and surfactant release into airspaces (48, 49, 72–75). Although reports indicate that extreme stretch of alveolar epithelial cells decreases surfactant secretion (76), we do not identify published data that show the effect of a stretch stimulus that is at a physiological level, but prolonged. In this regard, it is noteworthy that our findings show surfactant secretion was delayed in alveoli that were exposed to 15 min of static stretch by hyperinflation or hypoperfusion. Resolution of the stretch by reducing airway pressure or restoring lung perfusion rapidly stimulated surfactant release. We interpret from our findings that while alveolar stretch stimulated a surfactant secretion response in the alveolar epithelium, surfactant release occurred only after the stretch stimulus resolved. It seems physiologically appropriate that surfactant release would occur at the same time that lung deflation decreases the alveolar radius of curvature, since decrease of alveolar radius of curvature would raise surface tension at the airspace air-liquid interface. The mechanism that underlies the surfactant release delay is not yet clear. One possibility is that cytosolic Ca^2+^ responses that initiate surfactant secretion occur during the cell stretch phase of a transient stretch stimulus, but surfactant release from lamellar bodies occurs during the relaxation phase. If this is the case, the mechanism of surfactant release could relate to relaxation-induced cytoskeletal reorganization in AT2 cells that permits lamellar body fusion with AT2 cell plasma membranes. Published data support the notion that cytoskeletal reorientation occurs rapidly during cell relaxation (77). Whatever the mechanism, our findings show alveolar stretch by lung hyperinflation or hypoperfusion not only blocks AWL secretion, it also blocks surfactant secretion. Release of the stretch stimulus initiates secretion of both AWL and surfactant, indicating that AWL and surfactant secretion remain coupled in stretched alveoli of hypoperfused lungs.

Our findings may have clinical implications. Since AWL secretion contributes to alveolar defense by clearing particles and bacteria from alveolar walls (25, 26, 30) and providing a medium for alveolar macrophage function and surfactant and antibacterial peptide activity, loss of AWL and surfactant secretion in alveoli of hypoperfused lungs could promote lung infection. Hence, loss of AWL and surfactant secretion could be a mechanism by which acute lung diseases that are characterized by hypoperfusion – for example, acute chest syndrome, severe viral lung infection, ventilator-induced lung injury, and acute respiratory distress syndrome – lead to bacterial pneumonia. Pulmonary edema might further promote lung infection by causing expansion and stretch of neighboring, air-filled alveoli (78) that further impairs AWL– and surfactant-mediated alveolar defense. Over longer periods of time, hypoperfusion-induced inhibition of AWL secretion in upper lung regions, where West has identified that the ratio of ventilation to perfusion is high (79), could promote particle or pathogen deposition in upper lung alveoli and thus contribute to the pathogenesis of upper lung-predominant lung diseases like emphysema, tuberculosis, and hypersensitivity pneumonitis. Future research might define the extent to which the mechanisms we identified in this study contribute to the pathogenesis of human lung diseases.

Our study has some limitations. First, our findings were generated in mouse lungs and require confirmation in human lungs, since ion transport proteins can differ across species (80). Second, we used drug inhibitors that might have off-target effects. Although replicating our data using genetic inhibition approaches would be the ideal next step, achieving genetic inhibition of critical ion transport proteins is challenging due to problems with animal model viability and compensatory responses by other ion transport proteins. Inducible knockout models like those we have used for CFTR (63) may be helpful. We note VU0463271 seems to be KCC-specific (41, 42). Third, our intact, perfused lung model is roller pump-perfused and lacks the pulsatile flow of the circulatory system in vivo. This may be a consideration for follow-up studies since perfusion pulsatility impacts the regional distribution of blood flow in isolated dog lungs (81) and may affect the rate of lung fluid filtration (82). Finally, our preparation lacks ventilation-induced oscillatory changes of airway pressure. Although positive airway pressure has no effect on lung fluid balance in awake sheep (83), future research that tests the effect of oscillatory changes of airway pressure on alveolar liquid dynamics might lead to a fuller understanding of how lung hypoperfusion stimulates alveolar liquid absorption. Despite these limitations, our study provides strong that perfusion regulates alveolar liquid transport in lungs in which airway and perfusion pressures are tightly controlled.

In conclusion, our findings show lung hypoperfusion caused the alveolar epithelium to reverse liquid transport from homeostatic AWL secretion to airspace liquid absorption. AWL secretion was CFTR-, NKCC, and Na^+^/K^+^-ATPase-dependent, in line with known mechanisms. The switch to absorption occurred within minutes of lung hypoperfusion and was ENaC-, CFTR-, and KCC-dependent, and it was accompanied by inhibition of surfactant release from AT2 cells. We propose lung hypoperfusion caused alveolar liquid absorption by reducing microvascular lumen caliber, leading to alveolar epithelial stretch that might have inhibited NKCCs and activated KCCs. Together, these findings reveal a new role for KCCs in the lung, and they contribute new understanding of how alveolar function changes in response to hypoperfusion stress. Hypoperfusion-induced loss of liquid and surfactant secretion in alveoli might have a role in the pathogenesis of lung diseases characterized by acute microvascular hypoperfusion.

## METHODS

### Experimental design

The experiments were designed and the manuscript was written according to ARRIVE guidelines. Sample sizes and outcome measures are indicated in figures and legends.

### Fluorophores

We purchased calcein AM (final concentration 10 μM), calcein red-orange AM (10 μM), tetramethylrhodamine-conjugated dextran (TR-dextran, 70 kDa), fluorescein isothiocyanate-conjugated dextran (FITC-dextran, 20 kDa), lysotracker green (100 nM), and lysotracker red (100 nM) from ThermoFisher Scientific. Phycoerythrin (PE)-conjugated anti-CD11c antibody (8 µg/mL) was purchased from ThermoFisher Scientific.

### Reagents

All reagents were freshly constituted for experiments. We purchased HEPES and Ca^2+^– and Mg^2+^-containing PBS from ThermoFisher Scientific and NaCl, KCl, glucose, CaCl_2_, MgCl_2_, and sucrose from Sigma. We purchased CFTRinh172 (20 μM), bumetanide (1 μM), ouabain (1 mM), and VU0463271 (1 μM) from Sigma and amiloride (10 μM) from Selleck Chemicals. Tamoxifen was purchased from Sigma, reconstituted at 20 mg/mL in sunflower oil (Sigma), and administered within 30 min of reconstitution.

### Solutions

Calculated and measured values for solution contents are detailed in **Supplemental Table 1** and compared with reported normal values for mouse blood pH (84), plasma osmolality (36), and plasma Na^+^, K^+^, Cl^-^, ionized Ca^2+^, and non-fasting glucose (85–87) and reported values for plasma osmolality in dehydrated mice (36, 37). Normal values for the alveolar lining liquid are not known, as far as we are aware, for unchallenged adult animals or humans. We prepared HEPES-buffered solution to serve as the control instillation solution and as the vehicle for the lung perfusate and all airspace, airway, and nebulizer instillations. Calculated values for the HEPES-buffered solution are Na^+^ 150 mM, K^+^ 5 mM, Ca^2+^ 1 mM, Cl^-^ 159 mM, HEPES 20 mM, and glucose 10 mM and as otherwise indicated in **Supplemental Table 1**. A pH meter (Mettler Toledo Seven Excellence) and NaOH were used to titrate the solution pH. Osmolality was measured by freezing point depression using a micro-osmometer (Precision Systems 6002 Touch Micro Osmette). To generate the lung perfusate solution, we modified the HEPES-buffered vehicle solution by adding dextran (70 kDa; Molecular Probes; 4% w/v) and FBS (Gemini Bio-Products; 1% w/v). In separate experiments, we further added sucrose to generate the hyperosmolar lung perfusate solution, FITC-dextran (5 mg/mL) for alveolar barrier determinations, and drug inhibitors to determine the role of ion transport proteins in alveolar liquid dynamics.

### Animals

Wild type mice were C57BL/6J and 10-12 weeks old at the time of imaging. Transgenic mice were on a C57BL/6J background and 12-36 weeks old. Transgenic strains expressed tamoxifen-inducible, *Sftpc-CreER^T2^* (88) (Jackson Laboratory strain #028054), tamoxifen-inducible, *Ager-CreER^T2^* (89) (Jackson Laboratory strain #032771), and *ROSA^mT/mG^* (90) (Jackson Laboratory strain #007676) as indicated in the text and figure legends. All Cre-expressing mice were genotyped by Transnetyx using tail biopsy performed within 2 weeks of age. To induce *Cre* expression, we gave intraperitoneal injections of 100 μL of tamoxifen in sunflower oil at 8 weeks of age.

### Sex as a biological variable

Our study examined primarily male mice in order to achieve a high level of standardization for our experiments and minimize experimental variability from sources that were not directly related to perfusion of the isolated lung preparation. Data generated using female mice are indicated in **Supplemental Figure 3**.

### Bacterial strain and preparation

*S. aureus* (SA) was GFP-expressing USA300 LAC. SA were stored at –80°C in 25% glycerol in autoclaved Luria-Bertani (LB) broth media (MP Biomedicals) and propagated on LB-agar plates containing chloramphenicol (10 ug/mL). Plates were refreshed from frozen stock every 2 weeks. Single colonies of SA were propagated in autoclaved LB media containing chloramphenicol (10 μg/mL) in a shaking incubator at 37°C and 200 rpm (New Brunswick Scientific) for 18 h (stationary growth phase), then the bacterial solution was prepared for intranasal instillation by diluting 1.3 mL in 200 uL of PBS containing Ca^2+^ and Mg^2+^.

### Intranasal instillation

SA were prepared for intranasal instillation and instilled within 10 min of bacterial removal from the incubator. Mice were anesthetized with inhaled isoflurane (4%) and intraperitoneal injections of ketamine (up to 1 mg) and xylazine (up to 0.1 mg). Each mouse was instilled with 30 μL of prepared SA solution to deliver 1 x 10^8^ CFU per mouse. Instillation quality was recorded at the time of instillation by the performing investigator and considered acceptable for experiments if no loss of instillate was observed. Mice woke from anesthesia within 3 min of instillation. At 5 h after intranasal instillation, mice were re-anesthetized with inhaled isoflurane (4%) and intraperitoneal injections of ketamine (up to 100 mg/kg) and xylazine (up to 5 mg/kg) for euthanasia and surgical preparation of isolated, perfused lungs.

### Nebulizer instillation

Mice were anesthetized with inhaled isoflurane (4%) and intraperitoneal injections of ketamine (up to 50 mg/kg) and xylazine (up to 2.5 mg/kg). Within 5 min of starting anesthesia, mice were removed from the anesthesia chamber and fitted with a nose cone connected to a nebulizer machine. We added 3 mL of TR-dextran (1.25 mg/mL in HEPES-buffered vehicle solution) to the nebulizer chamber and applied nebulization for 5 min. Mice were anesthetized but breathing spontaneously for the duration of the nebulization procedure. After nebulization, the mouse was given a second intraperitoneal injection of ketamine (for total dose up to 100 mg/kg) and xylazine (for total dose up to 5 mg/kg) and moved back to the anesthesia chamber for additional exposure to inhaled isofluorane (4%) to achieve euthanasia for surgical preparation of isolated, perfused lungs for imaging.

### Preparation of isolated, perfused lungs for imaging

We anesthetized mice with inhaled isoflurane (4%) and intraperitoneal injections of ketamine (up to 100 mg/kg) and xylazine (up to 5 mg/kg), then gave intracardiac injections of heparin (50 units; Mylan) and exsanguinated the mice by cardiac puncture. Using our reported methods (25, 26), we cannulated the trachea, pulmonary artery, and left atrium of the heart, then excised the heart, lungs, and cannulas en bloc. The lungs were positioned to enable micropuncture and imaging of the diaphragmatic surface of the right middle lobe, right caudal lobe, or left lung. Then, the lungs were inflated with room air through the tracheal cannula and perfused through the pulmonary arterial and left atrial cannulas with autologous blood diluted in the lung perfusate solution or the hyperosmolar lung perfusate solution (see **Supplemental Table 1**) and warmed to 37°C. We used in-line pressure transducers (ADInstruments) to maintain constant airway pressure 6 cm H_2_O via a continuous positive airway pressure (CPAP) machine (Philips Respironics) and pulmonary artery and left atrial pressures via a roller pump (Ismatec). See text for details regarding the position of the vascular cannulas. The roller pump rate was initiated at 250 μL/min, then increased in 250 μL/min increments over 10 min to achieve pulmonary artery pressure (PAP) 10 cm H_2_O. After 5 min of lung perfusion at PAP 10 cm H_2_O, the roller pump rate was adjusted in 250 μL/min increments to achieve the goal PAPs indicated in **Supplementary Table 2**. Perfusion was halted by stopping the roller pump. For all experiments, the left atrial pressure (LAP) cannula was at the height of the lung and open to the atmosphere. LAP in lungs perfused at PAP in the physiological range was similar to LAP reported in rat lungs by direct LAP measures (91). Portions of the lung surface that were not used for micropuncture and imaging were covered with plastic wrap to prevent desiccation.

### Lung hyperinflation

After establishing initial airway pressure 6 cm H_2_O via CPAP, we increased the airway pressure to 12 or 16 cm H_2_O using an air syringe fitted in line with the CPAP machine. Airway pressures were monitored continuously using our in-line pressure transducer and maintained at the goal pressure for the duration of the hyperinflation experiments.

### Airway instillation of drug inhibitors

To minimize alveolar exposure to micropuncture, prior to initiating lung perfusion we exposed the alveolar epithelium of some lungs to vehicle solution, CFTRinh172, or amiloride by instilling the trachea cannula with 100 μL of solution containing either HEPES-buffered vehicle solution plus TR-dextran, or HEPES-buffered vehicle solution plus TR-dextran plus CFTRinh172 or amiloride. After the instillation we initiated lung perfusion as indicated in *Preparation of isolated, perfused lungs for imaging*. After achieving goal PAP, we instilled alveolar airspaces with calcein AM, then TR-dextran (10 mg/mL) by micropuncture (see *Alveolar microinstillation*).

### Alveolar microinstillation

We hand-beveled glass micropipettes (Sutter Instruments) to micropuncture single alveoli under bright-field microscopy, as we have done previously (25, 26). Micropunctured alveoli were instilled with fluorophores and reagents in solution, resulting in their spread from the micropunctured alveolus to neighboring alveoli. See text. Microinstillations were performed in 1-3 alveoli bordering imaging fields.

### Live lung imaging and analysis

By our established methods (25, 26), we viewed alveoli by confocal microscopy (LSM800; Zeiss) with a 20x water immersion objective (NA 1.0; Zeiss) and coverslip. We used bright-field microscopy to randomly select regions of 30-50 alveoli for microinstillation and imaging. All images were acquired as single images using Zen (v.2.6; Zeiss) and recorded as Z-sections. Analyzed images were 4-8 μm below the pleura for most experiments. Optical thickness was 2 μm, and frame size was 512 x 512 pixels. We established laser, filter, pinhole, and detector settings at the beginning of each imaging experiment to optimize alveolar fluorescence, then maintained the settings for the duration of the experiment. We confirmed absence of bleed-through between fluorescence emission channels. Images were analyzed using ImageJ (NIH; v.2.0.0-rc-69/1.52n). Linear adjustments of brightness and contrast were applied to individual color channels of entire images and equally to all experiment groups. We did not apply downstream processing or averaging.

### Determinations of alveolar liquid transport

We used alveolar micropuncture (see *Alveolar microinstillation*) to microinstill alveolar airspaces with TR-dextran in HEPES-buffered vehicle solution as described in *Solutions* and **Supplemental Table 1**. We used ImageJ (NIH; v.2.0.0-rc-69/1.52n) to quantify dextran gray levels in alveolar airspaces. To account for minor heterogeneity of dextran fluorescence changes across imaged alveolar regions, dextran gray levels were quantified as a mean of dextran fluorescence in representative alveoli located at the image top left, top right, bottom left, bottom right, and center. Measurement locations were the same across time points. All experiments were followed by determinations of alveolar barrier permeability (see *Determinations of alveolar barrier permeability*). We excluded imaged regions of alveoli from the dextran analysis if initial dextran gray levels were less than 20 and in the rare event that permeability determinations showed leak of microvascular fluid into alveolar airspaces.

### Determinations of alveolar barrier permeability

To determine alveolar barrier properties, we added FITC-dextran (20 kDa; 5 mg/mL) to the lung perfusate solution using our established methods (25, 26). FITC-dextran was circulated in the perfusate for 10 min prior to image acquisition.

### Determinations of surfactant secretion

We induced surfactant secretion in alveoli pretreated with alveolar microinstillation of lysotracker green or lysotracker red by either halting lung perfusion for 10 min, then restarting it to achieve goal PAP 10 cm H_2_O, or by increasing lung airway pressure from 6 to 12 cm H_2_O for 15 min or 15 s, then returning to 6 cm H_2_O. Lysotracker fluorescence was quantified using maximum intensity Z projections of stacks of 6 images that encompassed whole AT2 cells and extended from approximately 2-12 µm below the pleura.

### Statistics

Statistics are indicated in figures and legends. In general, paired comparisons were analyzed using two-tailed *t* tests, and multiple comparisons were made using ANOVA with post hoc Tukey testing. We considered statistical significance at *p* < 0.05. Data were analyzed and figures were prepared using Microsoft Excel, StatPlus:mac Pro (AnalystSoft, Inc., Build 7.5.0.0/Core v7.6.11), and SigmaPlot (Systat, version 14.5).

### Study approval

The Institutional Animal Care and Use Committee of the Icahn School of Medicine at Mount Sinai approved the animal procedures.

### Data availability

All data are available in the manuscript, supplemental materials, or supporting data values.

## AUTHOR CONTRIBUTIONS

J.Z., D.C., and J.L.H. designed the study. All authors contributed to data collection, analysis, and interpretation or results. J.H. was responsible for the overall project and wrote the first draft of the manuscript. All authors edited the manuscript. The order of the co-first authors was determined based on J.Z.’s greater contribution to the project’s conceptual design.

## ACKNOWLEDGEMENTS

This work was supported by NIH grants R01HL164821 (J.H.) and F31HL179970 (S.T.), Cystic Fibrosis Foundation Research Grants 004792G222 and 009144G225 (J.L.H.), and a Fellowship from the Stony Wold-Herbert Fund, Inc. (J.Z.). We thank Drs. Jahar Bhattacharya and Sunita Bhattacharya for thoughtful critique and helpful suggestions.

## SUPPLEMENTAL DATA

**Supplemental Table 1.** Composition of airspace and lung perfusate solutions. Hyperosmolar lung perfusate solution was made by adding sucrose, as indicated, to the lung perfusate solution. Ionized Ca^2+^ values are corrected for pH. *D70*, dextran (70 kDa); *FBS*, fetal bovine serum. Citations for normal values are indicated in the Methods.

|  | Direct measures |  |  |  |  |  |  |  | Calculated values |  |  |  |
| --- | --- | --- | --- | --- | --- | --- | --- | --- | --- | --- | --- | --- |
|  | pH | osmolality<br>(mOsm/L) | Na<sup>+</sup><br>(mM) | K<sup>+</sup><br>(mM) | Cl<sup>-</sup><br>(mM) | ionized<br>Ca<sup>2+</sup><br>(mM) | glucose<br>(mg/dL) | sucrose<br>(mM) | Mg<sup>2+</sup><br>(mM) | HEPES<br>(mM) | D70<br>(% w/v) | FBS<br>(% w/v) |
| Normal blood and plasma values in mice | 7.4 | 323-330 | 146-151 | 3.8-4.4 | 107-111 | 1.10-1.13 | 164-176 |  |  |  |  |  |
| Blood values in mice exposed to dehydration |  | 336-343 |  |  |  |  |  |  |  |  |  |  |
| Lung perfusate solution | 7.4 | 333 | 153 | 4.9 | 143 | 1.09 | 171 |  | 1.0 | 20 | 4 | 1 |
| Hyperosmolar lung perfusate solution |  | 360 |  |  |  |  |  | 20 |  |  |  |  |
| HEPES-buffered vehicle solution for instillations | 7.4 | 324 | 151 | 4.8 | 143 | 1.00 | 168 |  | 1.0 | 20 | 0 | 0 |
| plus TR-dextran at 10 mg/mL |  | 361 |  |  |  |  |  |  |  |  |  |  |
| plus TR-dextran at 1.25 mg/mL |  | 331 |  |  |  |  |  |  |  |  |  |  |

**Supplemental Table 2.** Lung perfusion pressures. Pulmonary artery pressure (*PAP*) and left atrial pressure (*LAP*) were measured in intact, perfused lungs of at least 3 mice using in-line pressure transducers located at the pulmonary artery and left atrial cannulas. See text. Citations for normal values are included in the Results for mouse PAP, and in the Methods for rat LAP. *Note, literature for rat LAP is used rather than mouse in order to provide data generated by direct LAP measurements.

| PAP group | PAP (cm H<sub>2</sub>O) | LAP (cm H<sub>2</sub>O) |
| --- | --- | --- |
| Normal values | 13-25 | 4* |
| PAP 15 | 15.7 ± 3.7 | 3.7 ± 0.3 |
| PAP 10 | 10.6 ± 2.3 | 2.3 ± 0.1 |
| PAP 5 | 5.5 ± 0.1 | 1.1 ± 0.1 |
| PAP 0 | 2.5 ± 0.0 | 0.0 ± 0.0 |

**Supplemental Figure 1.**
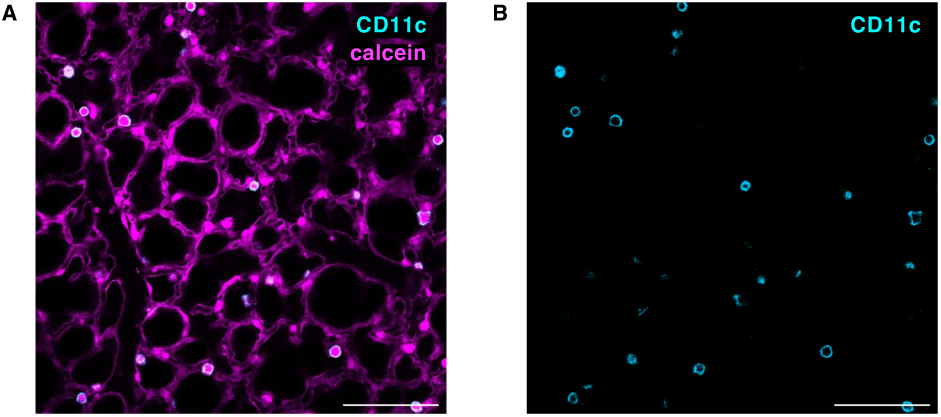
Innate immune cells in live alveoli of intact, perfused lungs. (**A-B**) Confocal images show CD11c+ cells (*cyan*) in live alveoli of mouse lungs. Alveolar walls are delineated by fluorescence of calcein (*magenta*). To generate the images, we microinstilled alveolar airspaces with phycoerthrin (PE)-tagged anti-CD11c antibody, then HEPES-buffered vehicle solution to wash out free antibody from airspaces. B is the same image as A, but calcein fluorescence has been digitally removed to better demonstrate CD11c fluorescence. The example images were replicated in lungs of 3 mice. Scale bar: 100 µm.

**Supplemental Figure 2.**
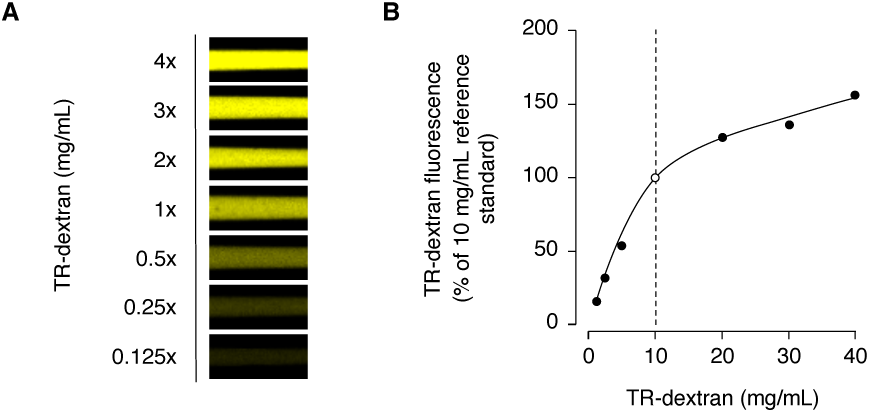
Fluorescence of TR-dextran in glass micropipettes. (**A-B**) Confocal images (A) and plot (B) show the relationship between concentration of tetramethylrhodamine-conjugated dextran (*TR-dextran*, 70 kDa, *yellow*) and fluorescence intensity in aqueous solution in glass micropipettes. The *open circle* and *dashed line* in B indicate the 10 mg/mL reference standard. The *solid line* was calculated by polynomial regression.

**Supplemental Figure 3.**
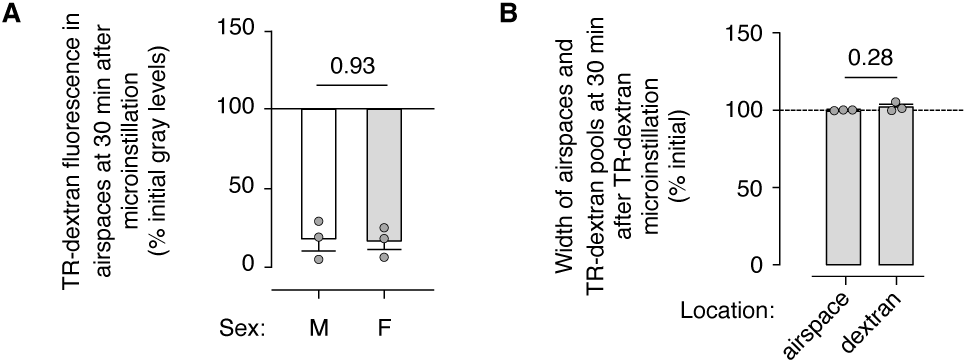
AWL secretion in lungs of male and female mice. (**A-B**) Confocal imaging experiments were carried out as in Figure 1, A-B. Perfusion was maintained at PAP 10 cm H_2_O. The data representing experiments in male mice (*M*) are indicated by the *white bar* and are reproduced here from Figure 1F. All other data were generated in female (*F*) mice. The group data show change of alveolar TR-dextran fluorescence (A), TR-dextran pool width (B) and airspace width (B) over time. *Circles* indicate n and each represent 1 mouse in which mean TR-dextran fluorescence change was quantified in imaging fields of at least 50 alveoli (*M*) or 1 trial of TR-dextran microinstillation in which mean fluorescence change was quantified in an imaging field of at least 50 alveoli (*F*). In female mice, trials were carried out in lungs of at least 2 mice. *Bars*: mean ± SEM; *p* as indicated by two-tailed *t* test.

**Supplemental Figure 4.**
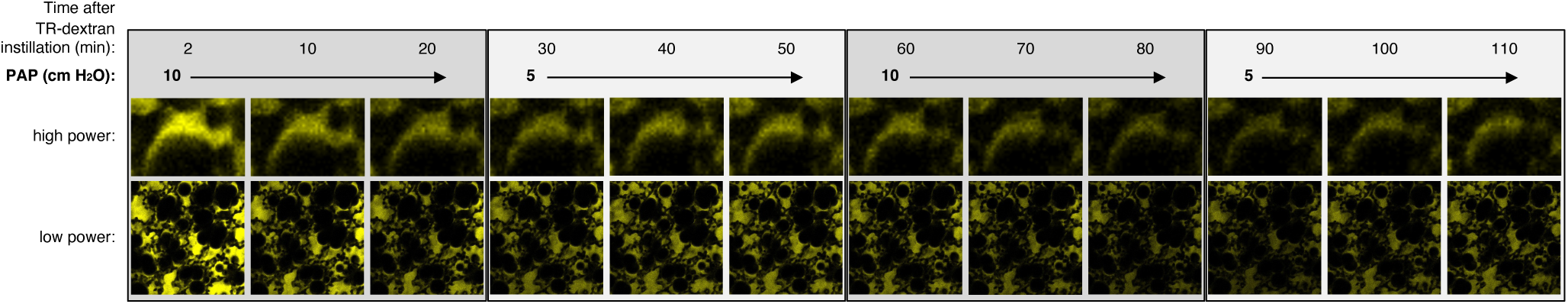
Alveoli respond rapidly to lung hypoperfusion: additional images. Confocal images are an expanded dataset of the example images shown in Figure 2C and show change of TR-dextran (70 kDa; *yellow*) fluorescence in alveoli of live, intact lungs perfused at the indicated pulmonary artery pressures (PAP). The example images were replicated in 3 trials of alveolar TR-dextran microinstillation in imaging fields of at least 50 alveoli. Scale bar is shown in Figure 2B.

**Supplemental Figure 5.**
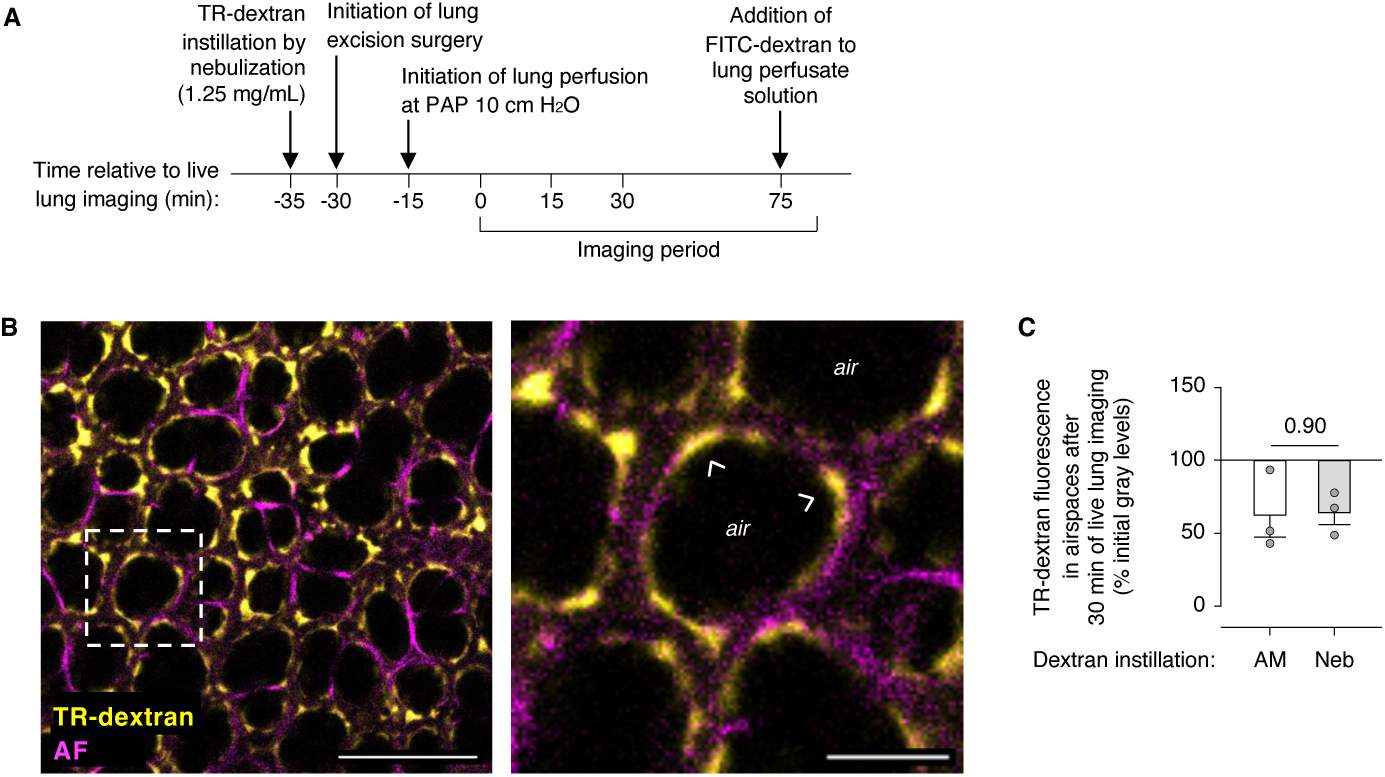
AWL secretion is not an effect of alveolar micropuncture. (**A-C**) Cartoon in <u>A</u> shows the design of experiments used to generate the confocal imaging data shown in B-C. As indicated in A, we exposed mice to TR-dextran (70 kDa) by nebulization for 5 min prior to lung excision for live lung imaging. The live lungs were perfused at PAP 10 cm H_2_O. In <u>B</u>, low (*left*) and high (*right*) power confocal images show TR-dextran (*yellow*) in alveoli in which alveolar walls are marked by autofluorescence (*AF*). *Arrowheads* point out where nebulized TR-dextran (*yellow*) pooled in alveolar niches. *Air,* example airspace. Group data in <u>C</u> show change of alveolar TR-dextran fluorescence after TR-dextran instillation by alveolar micropuncture (*AM*) or nebulization (*Neb*). Data represented by the *white bar* are reproduced from Figure 3C. In C, *circles* indicate n and each represent 1 mouse in which mean TR-dextran fluorescence change was quantified in imaging fields of at least 50 alveoli; *bars* show mean ± SEM; *p* as indicated by two-tailed *t* test. Scale bars: 100 (*left*) and 30 (*right*) µm.

**Supplemental Figure 6.**
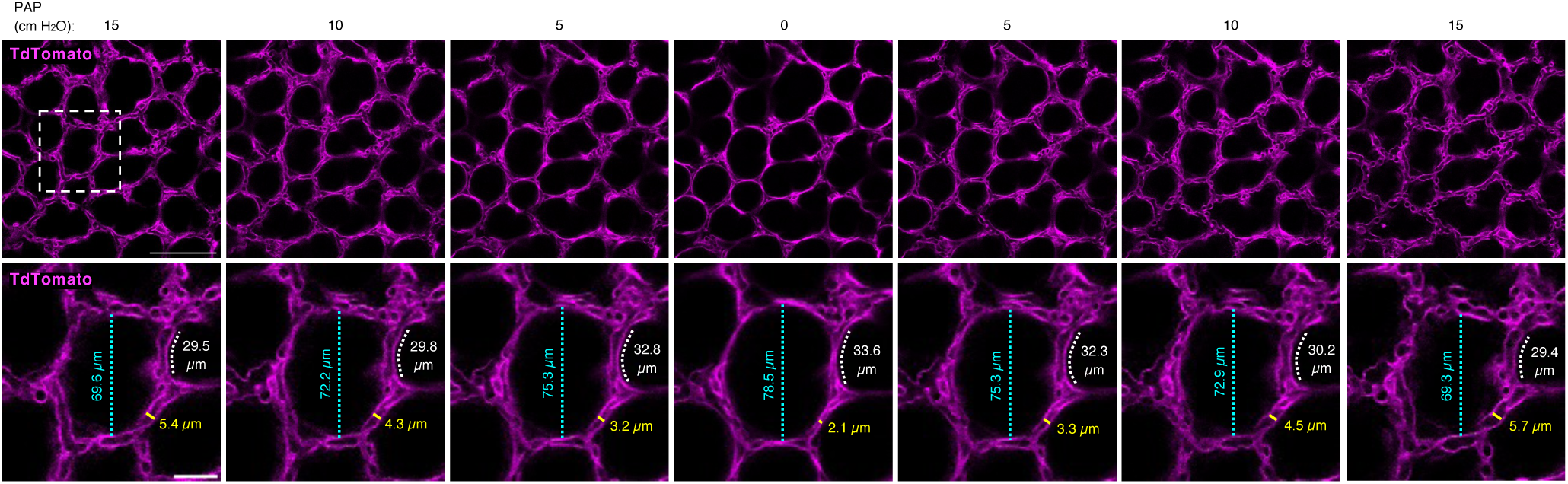
Lung hypoperfusion causes alveolar epithelial stretch: additional images. Confocal images are an expanded dataset of the example images shown in Figure 5B and show changes of alveolar morphology in response to changes of lung perfusion to the indicated pulmonary artery pressures (*PAP*). *Cyan lines and text* indicate alveolar airspace lumen diameter; *white lines and text* indicate alveolar septal length; and *yellow lines and text* indicate alveolar microvascular lumen diameter. Images were replicated in 4 imaging fields of at least 50 alveoli. Scale bars: 100 (*top row*) and 30 (*bottom row*) µm.

**Supplemental Figure 7.**
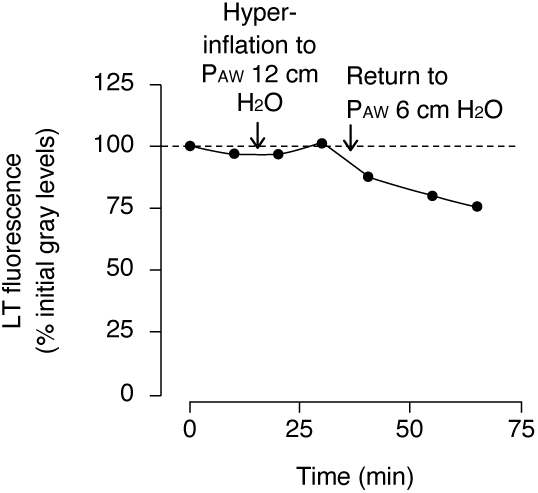
Surfactant secretion occurs after the release of sustained lung hyperinflation. The plot of confocal imaging data shows the effect of sustained lung hyperinflation on the fluorescence of AT2 cell lamellar bodies loaded with lysotracker dye (*LT*). Images were generated in isolated mouse lungs perfused at pulmonary artery pressure 10 cm H_2_O. As indicated, we initiated airway pressure (P_AW_) at 6 cm H_2_O, then increased P_AW_ to 12 cm H_2_O for 15 min, then decreased P_AW_ back to 6 cm H_2_O for the remainder of the experiment. Note, LT fluorescence is steady until after lung hyperinflation is released, when it decreases. LT fluorescence was quantified as mean gray levels in one imaging field that contained more than 20 AT2 cells.

